# Neurons that Function within an Integrator to Promote A Persistent Behavioral State in *Drosophila*

**DOI:** 10.1101/735985

**Authors:** Yonil Jung, Ann Kennedy, Hui Chiu, Farhan Mohammad, Adam Claridge-Chang, David J. Anderson

**Affiliations:** Division of Biology 156-29, Howard Hughes Medical Institute, TianQiao and Chrissy Chen Institute for Neuroscience, California Institute of Technology, Pasadena, CA 91125; Neuroscience & Behavioural Disorders Programme, Duke-NUS Medical School, Singapore, 138673; Institute of Molecular and Cell Biology, Singapore, 138673; College of Health & Life Sciences, Hamad Bin Khalifa University, Doha, Qatar

## Abstract

Innate behaviors involve both reflexive motor programs and internal states. In *Drosophila*, optogenetic activation of male-specific P1 interneurons triggers courtship song, as well as a persistent behavioral state that prolongs courtship and enhances aggressiveness. Here we identify pCd neurons as persistently activated by repeated P1 stimulation. pCd neurons are required for P1-evoked persistent courtship and aggression, as well as for normal social behavior. Activation of pCd neurons alone is inefficacious, but enhances and prolongs courtship or aggression promoted by female cues. Transient female exposure induced persistent increases in male aggressiveness, an effect suppressed by transiently silencing pCd neurons. Transient silencing of pCd also disrupted P1-induced persistent physiological activity, implying a requisite role in persistence. Finally, P1 activation of pCd neurons enhanced their responsiveness to cVA, an aggression-promoting pheromone. Thus, pCd neurons function within a circuit that integrates P1 input, to promote a persistent internal state that enhances multiple social behaviors.

## INTRODUCTION

Animal behaviors triggered by specific sensory cues evolve over multiple time-scales, from rapid reflex reactions to more enduring responses accompanied by changes in internal state^1, 2^. The former allow survival reactions, while the latter afford time to integrate contextual and other influences on behavioral decisions. In *Drosophila melanogaster*, male-specific P1 interneurons^3^ are activated by female-specific pheromones^4–6^, and control male courtship behaviors such as singing^7, 8^, as well as internal states that regulate aggression^9^, mating^10, 11^, feeding^12^ and sleep^13^ (reviewed in Ref. ^14^). Artificial stimulation of P1 neurons in solitary males can trigger rapid-onset courtship song^7, 8, 15^. Nevertheless, singing persists for minutes after stimulation offset^15, 16^, while optogenetically evoked P1 activity itself returns to baseline in tens of seconds^9, 15^ (but see ref ^17^). Similarly, the effect of P1 activation to promote aggressiveness endures for minutes after photostimulation offset^18^. These data suggest that persistent behavioral states evoked by P1 stimulation are not encoded in P1 neurons themselves, but rather in one or more of their downstream targets. We therefore sought to identify such targets, and to understand their functional role in the encoding of persistent behavioral states.

## RESULTS

To search for P1 follower cells exhibiting persistent responses, we expressed the red-shifted opsin Chrimson^19^ in P1^a^-split GAL4 neurons^9, 18, 20^, and a calcium indicator (GCaMP6s^21^) in ∼2,000 Fruitless (Fru)-LexA^22^ neurons (Fig. 1a). Optogenetic stimulation was calibrated to activate P1 cells at a level comparable to that evoked in these cells by female abdomen touching in the same preparation. Fru^+^ cells activated by P1 stimulation were identified by volumetric imaging (30 4-µm optical sections covering a 250 µm x 250 µm x 120 µm volume; Extended Data Fig. 1 d, e). On average, we monitored activity of 191 Fru^+^ cell somata and identified ∼37 cells per fly that responded to P1 stimulation (≥2/3 trials evoking a peak ΔF/F response >4σ above baseline; ED Fig. 1f), in 14 distinct brain regions. Different putative P1 follower cells showed different response durations, in a continuous distribution ranging from those similar to P1 (τ∼15 s; see Methods) to those lasting much longer (Fig. 1b, ED Fig. 1g, i). We used several criteria to select cells for further study: 1) median tau value > 5-fold that of P1 (τ>∼75 s); 2) persistent P1 responses detected in >75% of tested flies (n=12); 3) >2 cells/fly per hemibrain; 4) cells genetically accessible using specific GAL4 drivers.

**Figure 1.**
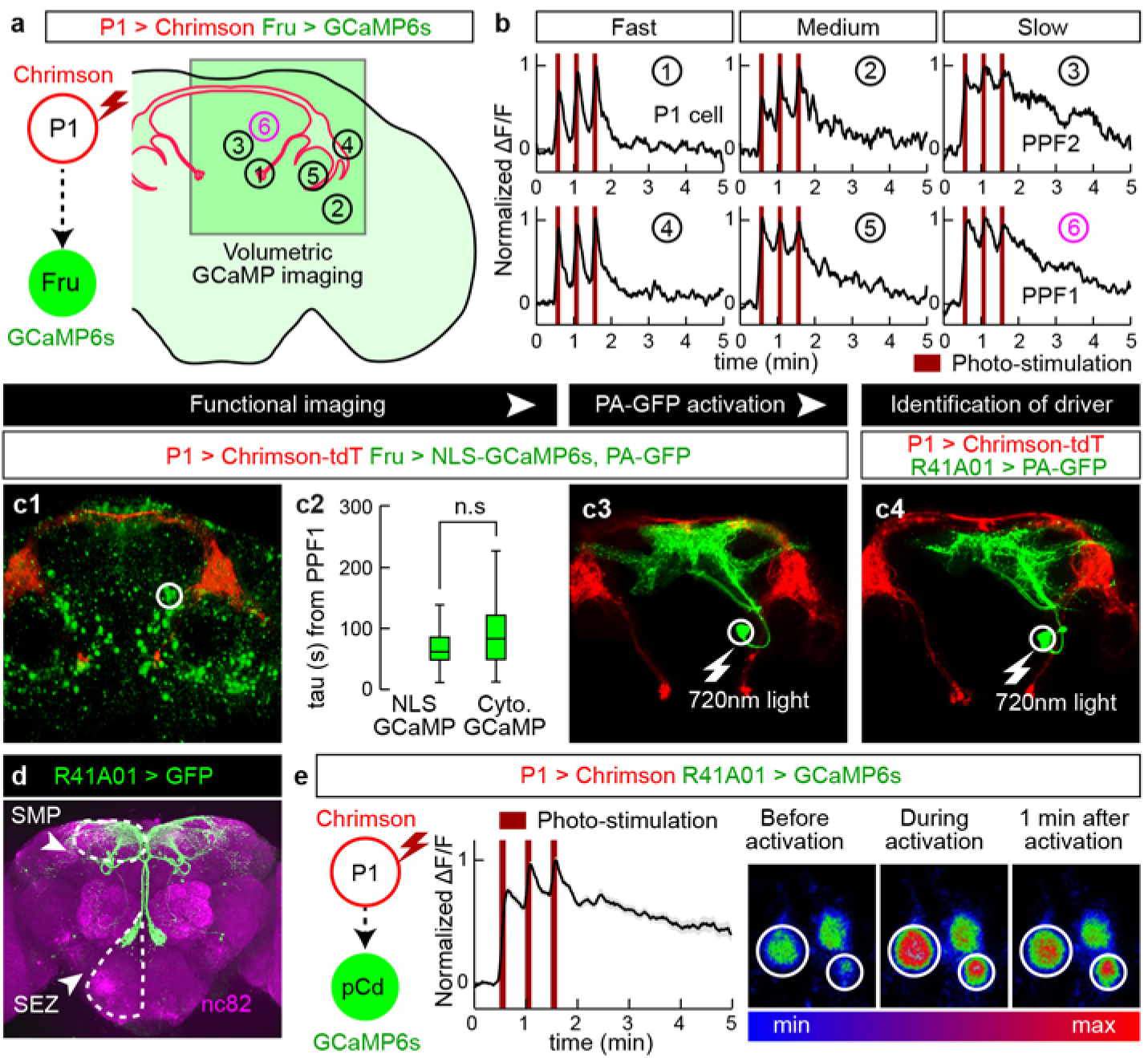
Identification of P1 follower cells with long-lasting responses. (a) Experimental schematic. Green square indicates imaging field containing different putative P1 follower cells (numbered circles). (b) Representative GCaMP6s traces (normalized ΔF/F); numbers correspond to cells in (a). PPF1 cells (➅) are pCd neurons. 655 nm light (10 Hz, 10 ms pulse-width, 25 s inter-stimulation interval) was delivered for Chrimson stimulation (dark red bars). (c1-4) Identification of GAL4 driver labeling PPF1 (pCd) neurons (See ED Fig. 2a for details). (c1) LexAop-NLS-GCaMP expressed in Fru-LexA neurons; white circle, PPF1 somata. (c2) Comparison between NLS GCaMP6s and Cytoplasmic GCaMP6s. Decay constants (tau) were calculated by curve fitting (See ED Fig. 1i and Methods for details). n=32 trials, 11 cells from 7 flies (NLS GCaMP), 77 cells from 12 flies (Cytoplasmic GCaMP). Statistical significance in this and in all other figures (unless otherwise indicated) was calculated using a Mann-Whitney U-test. Boxplots throughout show the median (center line), 25^th^ and 75^th^ percentiles (box), and 1.5 times the interquartile range (whiskers). Outliers were defined as data points falling outside 1.5x the interquartile range of the data, and were excluded from plots for clarity, but not from statistical analyses. (c3) PPF1 projections revealed by Fru-LexA>PA-GFP activation^23^. (c4) PPF1 neurons labeled by R41A01-LexA>PA-GFP. Non-PPF1 PA-GFP and NLS-GCaMP basal fluorescence have been masked for clarity. All images in c1, c2, and c4 are maximum intensity z-projections of 2-µm optical sections acquired by 2-P imaging. (d) Central brain R41A01 Gal4 neurons revealed by UAS-myr::GFP reporter. Superior medial protocerebrum (SMP) and sub-esophageal zone (SEZ) are indicated by dashed outlines. (e) LexAop-GCaMP6s response of pCd neurons labeled by R41A01-LexA following P1-Gal4/UAS-Chrimson stimulation (see Supp. Table 1 for genotypes). Left, schematic; middle, normalized ΔF/F trace (n=23 trials, 15 cells from 10 flies; mean±sem); right, fluorescent images taken before, during, and 1 minute after P1 activation (averaged over 5 frames). White circles indicate two responding cells.

We identified several putative persistent P1 follower (PPF) cells, which met the first criterion. These neurons were present in ∼5 distinct clusters, each containing ∼1-3 PPF cells, within a relatively small brain region (see Fig. 1a). Cells in one such cluster, PPF1 (Fig. 1b, #6) exhibited a median τ∼83 s (ED Fig. 1g, h). Cells in three other clusters including PPF2 (Fig. 1b, #3), showed a median τ>∼75, but failed to meet the second and third criteria. Another cluster in addition to PPF1 met all 3 criteria, but was not genetically accessible.

To gain specific genetic access to PPF1 neurons, we first examined the anatomy of these cells by combining P1 stimulation-evoked GCaMP imaging with photo-activatable GFP (PA-GFP) labeling of responding cells^23^. We generated a nuclear-localized GCaMP (NLS-GCaMP6s) to prevent cytoplasmic GCaMP signal from obscuring PA-GFP fluorescence (Fig. 1c1). NLS-GCaMP6s also detected persistent responses to P1 stimulation in PPF1 cells (Fig. 1c2). We then focused a 720 nm two-photon laser on the identified PPF1 cells, and revealed their projection pattern via diffusion of activated PA-GFP^23^ (Fig. 1c3). By comparing the morphology of PPF1 neurons with Fru-MARCM^24, 25^ and Gal4 line image databases^26^, we identified two Gal4 drivers, R41A01 and R21D06, which labeled morphologically similar neurons (Fig. 1c4, d; ED Fig. 2a-d). To verify that R41A01 and R21D06 indeed label PPF1 neurons, we performed functional imaging in R41A01>GCaMP6s or R21D06>GCaMP6s flies, and confirmed persistent responses to P1 activation in PPF1 somata (Fig. 1e and ED Fig. 2c); whether such persistent responses are present in all neurites is difficult to ascertain. Interestingly, these neurons exhibited stepwise integration of P1 input (Fig. 1e); however repeated P1 stimulation trials (as done in volume imaging, 30 trials, Fig. 1b) sensitized PPF1 neurons (ED Fig. 3).

**Figure 2.**
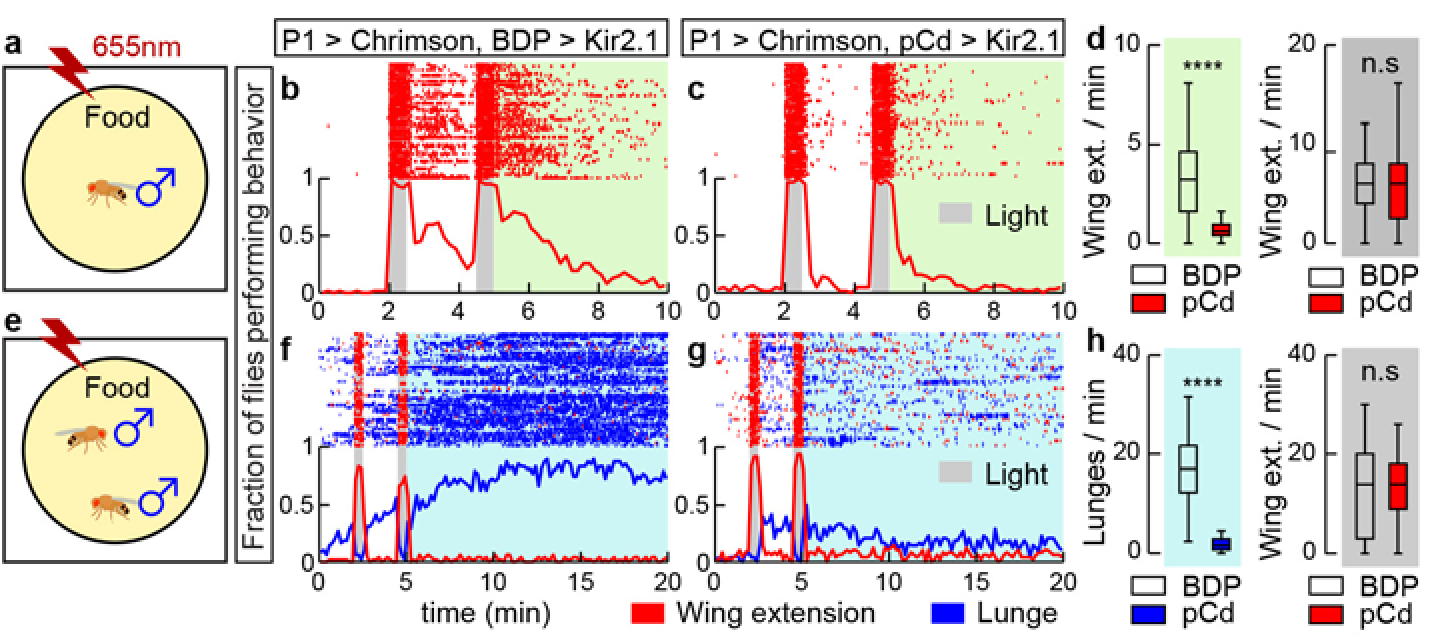
Activity of pCd neurons is required for P1-induced persistent behaviors. (a) Schematic (approximately to scale). Chrimson activation at 655 nm^15^ was performed in solitary males on food. (b-c) Behavior of flies during (gray shading) and after (green shading) P1^a9,20^ neuronal activation, either without (b; BDP is enhancerless LexA control driver), or with (c; pCd-LexA) Kir2.1-mediated^65^ inhibition of pCd neurons. Grey bars, 30 s photostimulations (40 Hz, 10 ms pulse-width) at 2 min intervals. Upper: Wing extension raster plot (red ticks). Lower: fraction of flies performing wing extensions (red line) in 10 s time bins. n=62 (b), 63 (c). (d) Wing extension frequency per fly after (green shading) or during (grey shading) photostimulation. **** *P* < 0.0001. (e) As in (a), but using male pairs. (f-g) Plot properties as in (b-c). Grey bars, 30 s photostimulation periods (2 Hz, 10 ms pulse-width) at 2 min intervals. Upper: raster plot showing wing extensions (red ticks) and lunges (blue ticks). Lower: fraction of flies performing wing extensions (red line) or lunges (blue line) in 20 s time bins. n=48 for each genotypes. (h) Lunge frequency after photostimulation (light blue shading, left), and wing extension frequency during photostimulation (grey shading, right). Lunging during, and wing extension after photostimulation were ≤ 1 event/min and are omitted for clarity. Statistics as in (d).

**Figure 3.**
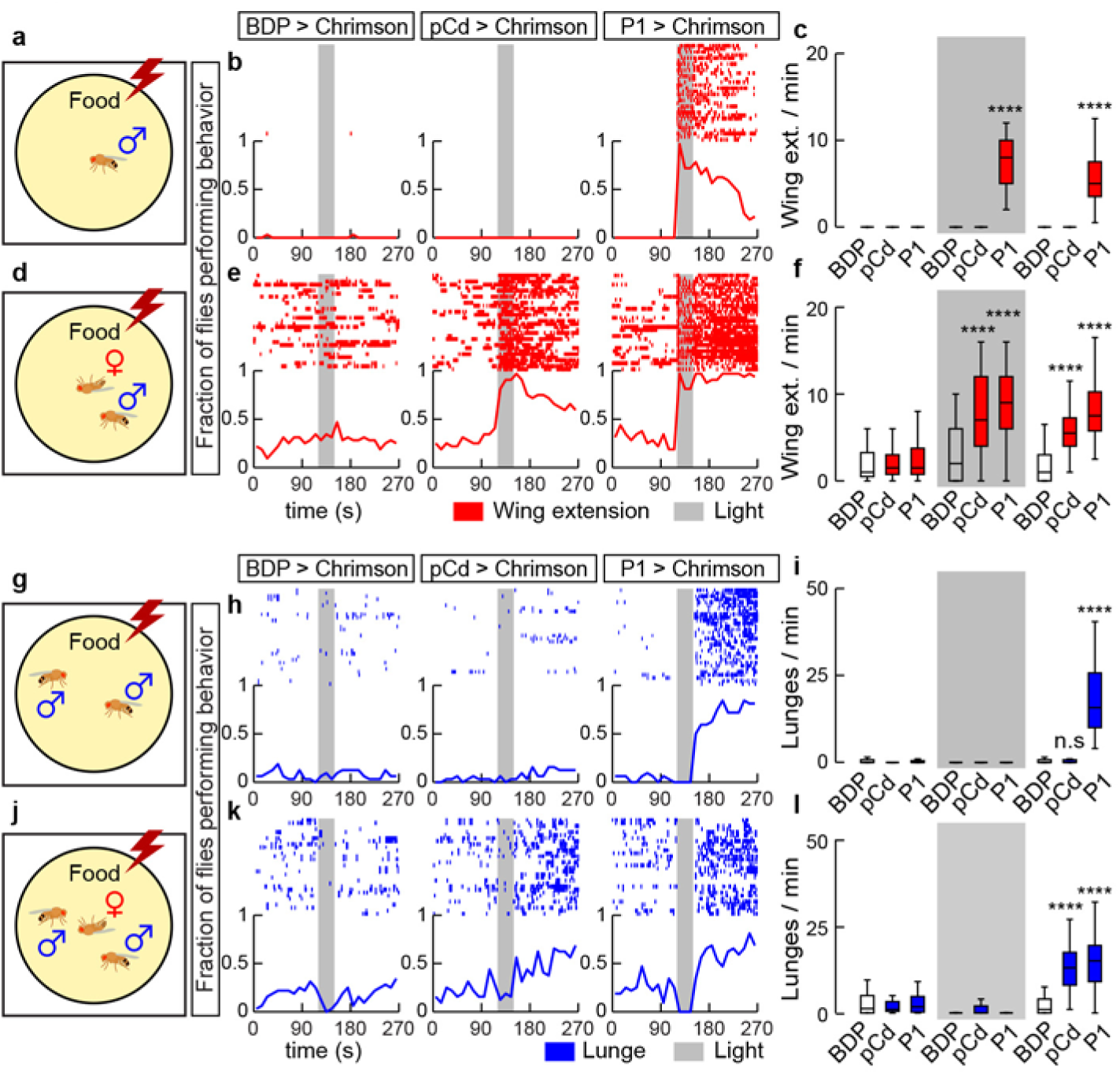
Activation of pCd neurons amplifies and extends male social behaviors induced by female cues. (a, d, g, and h), experimental schematics illustrating optogenetic activation of pCd neurons in solitary males (a-f) or pairs of group-housed males (g-l), tested without (a-c, g-i) or with (d-f, j-l) a dead female. Raster plots and fraction of flies performing behaviors (red and blue lines, 10 s time bins) are shown in (b, e, h, and k). Plot properties same as in Fig. 2. Grey bars, 30 s Chrimson activation at 655 nm (10 Hz, 10 ms pulse-width). Quantification and statistical tests shown in (c, f, i, and l). n=32 flies each. Statistical test used was a Kruskal-Wallis test. **** Dunn’s corrected *P* < 0.0001 for between-genotype comparisons. Courtship data are omitted in (h, k) for clarity.

Gal4 line R41A01 labels a cell cluster called pCd, previously reported to play an important role in female sexual receptivity^27^. Analysis of marker expression indicated that pCd cells are cholinergic neurons that express both Fru and Dsx (ED Fig. 2f-i), two sex-determination factors that label neurons involved in male courtship and aggression^3, 8, 28–31^. pCd neurons project densely to the superior-medial protocerebrum (SMP), while extending an additional long fiber bundle ventrally to innervate the dorsal region of the subesophageal zone (SEZ; Fig. 1d). Double labeling of pCd neurons with somatodendritic (Denmark-RFP^32^) and pre-synaptic (Syt-GFP^33^) markers revealed that their SMP projections are mostly dendritic, while their pre-synaptic terminals are located in the SMP and the SEZ (ED Fig. 4d-f). Registration of P1 pre-synaptic labeling with pCd somatodendritic labeling in a common brain template failed to reveal clear overlap (ED Fig. 4g-i), and application of the GFP Reconstitution Across Synaptic Partner (GRASP^34^) technique failed to detect close proximity between P1 and pCd neurons (ED Fig. 4j-r), suggesting that functional connectivity between these cells is unlikely to be monosynaptic.

**Figure 4.**
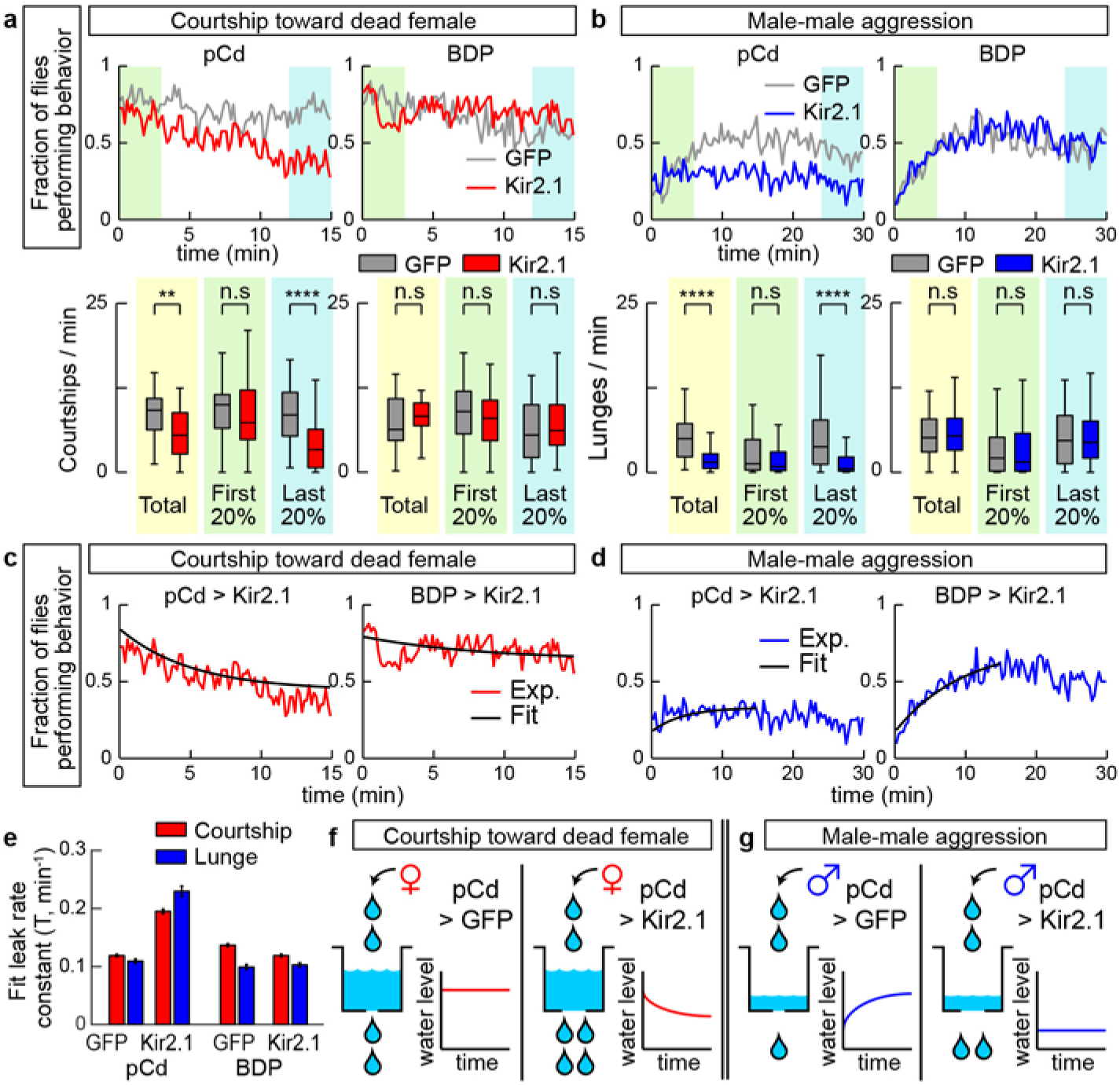
Inhibition of pCd neurons reduces endurance of naturally occurring social behaviors. (a) Solitary male flies were incubated with a dead female and courtship (unilateral wing extension bouts, UWEs) measured over 15 min. Left panels show experimental (pCd>Kir2.1, red line) and responder control (UAS-GFP, grey line) flies, right panels show enhancerless driver controls (BDP-Gal4; red and grey lines). Upper: fraction of flies performing behavior in 10 s time bins. Lower: number of UWE bouts per min per fly over entire 15 min observation (yellow shading), first (green shading) and last (blue shading) 20% (3 min) of the interval. n=40 flies per genotype. ** *P* < 0.01, **** *P* < 0.0001. (b) Pairs of single-housed males monitored over 30 min. Plot properties and statistical tests same as in (a), except blue color indicates lunging. Fraction of flies performing behavior was binned in 20 s time intervals. n=64 flies per genotypes. (c-d) Curve fitting of (c) courtship data from (a), or (d) lunging data from (b). Black lines show exponential fit curve for each experiment. Goodness of fit (MSE): courtship; 0.0042 (pCd>GFP), 0.0051 (pCd>Kir2.1), 0.0056 (BDP>GFP), 0.0058 (BDP>Kir2.1); Aggression; 0.0028 (pCd>GFP), 0.0031 (pCd>Kir2.1), 0.0045 (BDP>GFP), 0.0029 (BDP>Kir2.1). (e) Leak rate constants derived from curve fitting in (c, d); note that both courtship and lunging in pCd>Kir2.1 flies are best fit by assuming increased leak constants, relative to genetic controls. (f, g) Illustration of modeling results. Water level represents level of activity in a hypothetical leaky integrator driving behavior^40^. Inhibition of pCd activity with Kir2.1 increases leak rate constant of the integrator.

### pCd neuronal activity is required for P1-induced persistent social behaviors

To test whether P1-evoked persistent social behaviors require pCd activity, we silenced the latter using R41A01-LexA>LexAop-Kir2.1 while activating P1^a^-split GAL4 neurons using UAS-Chrimson. In solitary males (Fig. 2a), silencing pCd neurons dramatically reduced persistent wing extension evoked by Chrimson activation of P1 cells (Fig. 2b vs. 2c, green shading; 2d). Importantly, time-locked wing-extension during photostimulation was unaffected (Fig. 2b-d, gray shading). Persistent aggression evoked by P1 activation in pairs of males^9, 35^ (Fig. 2e, f) was also strongly reduced by silencing pCd neurons (Fig. 2g, h, blue shading), while wing-extension during photostimulation was unaffected. This result was confirmed using a more specific R41A01∩R21D06 intersectional split-GAL4 driver (ED Fig. 2d) to silence pCd neurons, and R15A01-LexA to activate P1 cells (ED Fig. 5). Thus pCd activity is required for enduring, but not for time-locked, behavioral responses to P1 activation.

**Figure 5.**
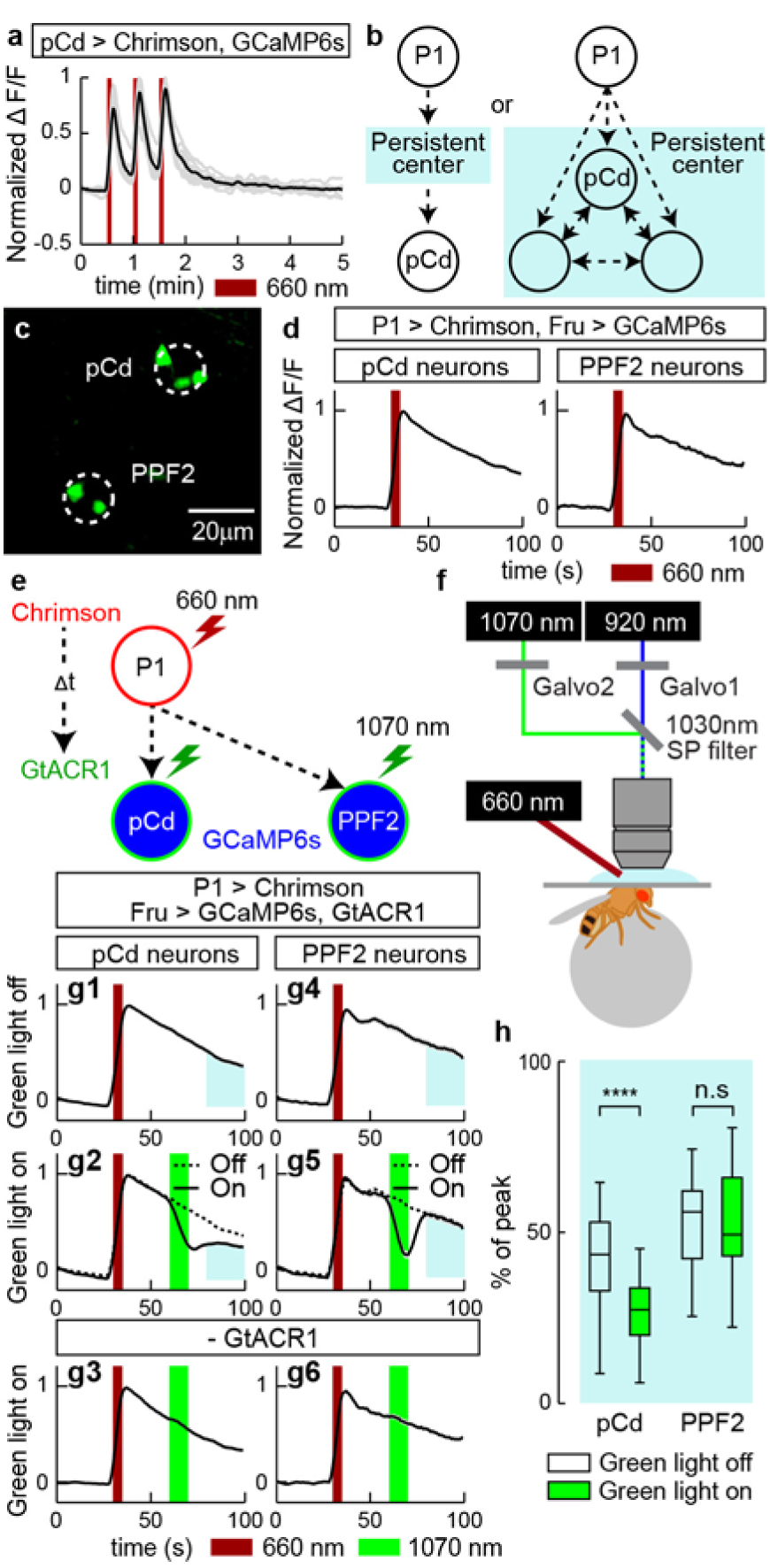
pCd neuronal activity is required for physiological persistence. (a) pCd response to direct optogenetic stimulation is not persistent. Gray lines depict individual pCd cell responses (n=27 from 9 flies), black line shows the mean for all cells. Dark red bars, Chrimson stimulation (655 nm light (10 Hz, 10 ms pulse-width, 25 s inter-stimulation interval). (b) Schematic illustrating alternatives tested by the experiment in (e-h). Light blue shading depicts hypothetical persistence-encoding network (“center”). If pCd neurons simply inherit persistence passively from the center (left), then persistence should rebound following transient pCd silencing. If persistence does not rebound, it implies that pCd activity is required for the center to maintain persistence (right). (c) Representative two-photon image showing cell body locations of pCd and PPF2 neurons expressing Fruitless>GCaMP6s in vivo. Dashed white circles indicate spiral scanning area for GtACR actuation in (e-h). Maximum intensity projection of 5 x 4 µm optical sections, averaged over 10 frames. (d) Normalized ΔF/F traces from pCd (left, n=36 trials from 8 flies), and PPF2 (right, n=29 trials from 5 flies) neurons upon P1 activation. Mean±sem. Dark red bar indicates P1 photostimulation (5 s, 10 Hz, 10 ms pulse-width, 660 nm LED). (e) Experimental schematic. pCd or PPF2 neuron cell bodies are locally photo-inhibited with GtACR1 (∼10 s, spiral scanning, see Methods for details) after a delay (Δt, 25 s) following P1 activation (5 s). (f) Schematic illustrating imaging setup with 1070 nm 2-photon laser for GtACR1 photo-inhibition, and 920 nm 2-photon laser for in vivo GCaMP imaging. (g) Normalized ΔF/F from pCd neurons (g1-g3), and PPF2 neurons (g4-g6) with GtACR actuation (green bars) applied during P1-induced persistent phase. g1 and g4: without photo-inhibition; g3 and g6, 1070 nm irradiation without GtACR1 expression. Dashed lines in g2 and g5 are mean of g1 and g4 traces, respectively. n=36 trials from 8 flies for pCd neurons, and 16 (5 flies) for PPF2 neurons. n=40 (8 pCd flies), 29 (6 PPF2 flies) for genetic controls. Mean±sem. (h) Normalized area under the curve (blue shaded regions in (g)) after photo-inhibition. **** *P* < 0.0001.

### pCd neurons amplify and prolong, but do not trigger, social behaviors

We next investigated the effect on behavior of optogenetically stimulating pCd neurons. Interestingly, optogenetic activation of pCd neurons in solitary flies had no visible effect, in contrast to optogenetic activation of P1 neurons^4, 9, 15^ (Fig. 3a, b). Persistent internal states can change an animal’s behavioral response to sensory cues. We reasoned that if pCd neurons promote such a persistent internal state, then their optogenetic activation, while insufficient to evoke behavior on its own, might nevertheless suffice to modify the behavioral response of the flies to an external social stimulus. To test this, we examined the effect of pCd stimulation on the behavioral response of males to female cues (Fig. 3d). Activation of pCd neurons in the presence of a dead female dramatically elevated courtship behavior during photostimulation, and this effect persisted for several minutes after stimulus offset (Fig. 3b vs. 3e, pCd>Chrimson; Fig. 3c, f, pCd).

Activation of pCd neurons in pairs of non-aggressive group-housed male flies did not promote aggression, unlike P1 activation^9^ (Fig. 3g-i). But in the presence of a dead female, which produced increased baseline aggression in male flies^36^, activation of pCd neurons significantly enhanced fly aggressiveness after photostimulation, an effect not observed in photostimulated controls (Fig. 3j-l). Thus, unlike P1 activation, which can substitute for the effect of dead females to trigger courtship or aggression, pCd activation alone cannot do so (Fig. 3b-c, h-i). However pCd neuron activation can enhance and extend the effect of a dead female to promote these social behaviors.

### pCd neurons are required for sustained courtship and aggressive drive

Given that pCd neuronal activity is required for optogenetic P1 activation-evoked social behavior (Fig. 2), we next investigated its requirement during natural social behavior. Silencing pCd neurons significantly increased the latency to copulation (ED Fig. 6a, b). To examine the effect of silencing on courtship *per se*, without rapid progression to copulation, we tested males in the presence of a freeze-killed virgin female, which induced robust unilateral wing-extensions (UWEs; courtship song^37^). In controls (BDP-GAL4> Kir2.1 or GFP), the fraction of flies exhibiting UWEs was relatively constant across the 15 min assay (Fig. 4a, BDP, gray and red lines). However in pCd>Kir2.1 flies, UWEs declined significantly during that interval, in comparison to pCd>GFP controls (Fig. 4a, pCd, red line, green vs blue shading).

**Figure 6.**
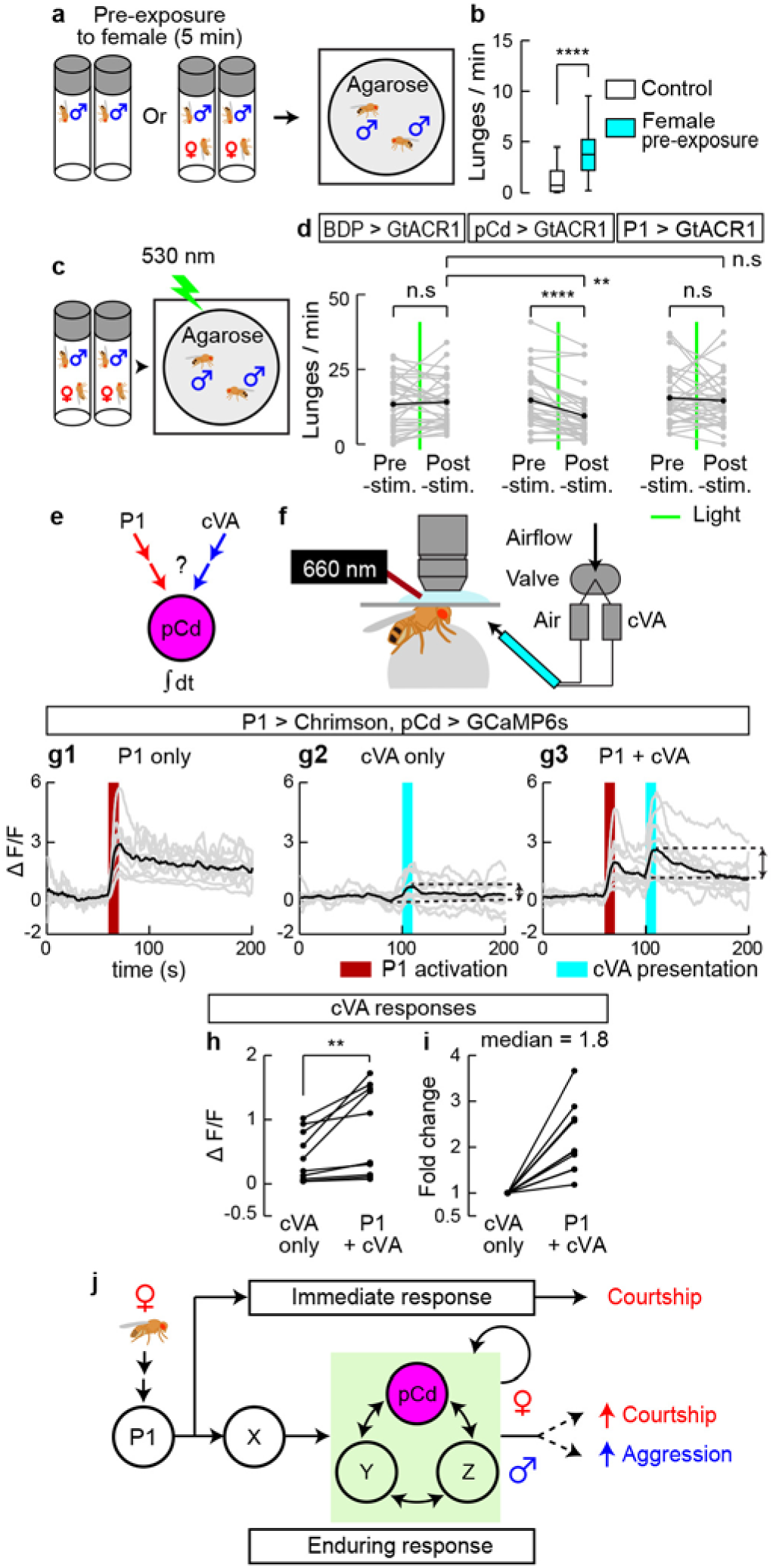
Role of pCd neurons in a female-induced enhancement of male aggressiveness. (a) Schematic illustrating female induced inter-male aggression experiment. Single-housed male flies were pre-incubated in vial with or without (control) a virgin female for 5 min. Subsequently, pairs of pre-incubated males were placed in behavioral arenas with an agarose substrate. (b) Lunge frequency per fly after pre-incubation without (white) or with (blue) a female. n=32 flies each. Statistical test used was a Mann-Whitney U-test. **** *P* < 0.001. (c) Schematic of experimental design. (d) Lunge number before (pre-stim.) and after (post-stim.) GtACR1-mediated neural silencing. Green lines depict exposure to green light (530 nm, 10 Hz, 10 ms pulse-width) for 10 s. Gray points show lunge frequencies for individual flies, and black points show mean values. Statistical tests used were Wilcoxon signed test (within fly comparison) and Kruskal-Wallis test (between genotype comparison). ** Dunn’s corrected *P* < 0.01, **** *P* < 0.0001 (e) Schematic illustrating experimental design. (f) In vivo GCaMP imaging. P1 neurons were optogenetically activated (660 nm LED), and cVA (or air) was delivered using an olfactometer synchronized and controlled by the imaging acquisition software. (g) GCaMP responses (ΔF/F) to cVA of pCd neurons exhibiting persistent responses to P1 photostimulation (g1, dark red bar, 10 s, 10 Hz, 10 ms pulse-width). cVA alone (g2, cyan bar) or 30 s after a second (10 s) P1 stimulation (g3) were delivered 3 min apart in random order (Methods). Gray lines depict trial-averaged individual pCd cell responses (2-3 trials/cell, n=10 cells from 7 flies) and black lines show the mean for all cells. Double-headed arrows in (g2 and g3) indicate intervals for cVA responses calculated in (h-i). (h) Individual pCd cell responses (ΔF/F) to cVA presented alone (“cVA only”) or 30 s after a 10 s P1 stimulation (“P1+cVA”). Statistical test used was a Wilcoxon signed-rank test. ** *P* < 0.01. (i) Fold change of pCd responses to cVA presentation after P1 stimulation, compared to cVA delivered alone. Data normalized to ΔF/F without P1 stimulation. (j) Models for how P1 and pCd neurons regulate immediate and enduring social behaviors.

We next performed parallel experiments for aggression. Single-housed (SH) male flies will fight on food in the absence of females^36, 38^, and the intensity of fighting escalates over time (Fig. 4b, BDP). However in SH pCd>Kir2.1 flies, aggression did not escalate over time, although initial levels of lunging were similar to controls (Fig. 4b, pCd, blue line, green vs. blue shading). These data demonstrate a requirement for pCd neurons in escalated aggression, independent of any influence from females. Importantly, in both assays, silencing pCd neurons did not impair initiation of social behavior, consistent with the inability of pCd optogenetic stimulation to trigger these behaviors (Fig. 3b, g); rather it influenced their amplitude and kinetics.

The effect of pCd silencing on courtship vs. aggression was subtly different: in the former case, silencing pCd neurons caused UWEs to steadily decline over time, whereas during aggression, natural escalation failed to occur (Fig. 4a vs. 4b, pCd, red vs. blue lines). To investigate whether a common mechanism could explain both phenotypes, we asked whether both data could be jointly fit by a “leaky integrator” model^39^. Such models formalize classical “hydraulic” theories of behavioral drive^40^, in which the instantaneous level of activity in a neural integrator circuit determines either the rate or type of an animal’s behavior; here, we sought to fit the time-evolving rate of UWEs, or of lunging (Fig. 4c). Our leaky integrator model assumed that flies received sensory input from conspecifics with a rate constant R, and that the activity of the neural circuit integrating conspecific sensory cues decayed from its initial condition to steady-state with a “leak” rate constant T (min^−1^).

The behavioral data in each assay were well fit by models in which the only free parameter allowed to vary by genotype was T (Fig. 4c-e). For UWEs, in control flies the relatively flat line reflects the fact that the initial rate of behavior is high, and already close to the steady-state where “fill” and “leak” rates are equal (Fig. 4f, left). In contrast, the faster decline of UWEs in pCd>Kir2.1 flies (Fig. 4c) was best fit by an increase in T (Fig. 4e, red bars). During aggression, control flies exhibit escalation (Fig. 4d, BDP>Kir2.1) because the initial rate of aggression is low, and the sensory input rate constant R is greater than T for this behavior (Fig. 4g, left). Increasing T in pCd>Kir2.1 flies therefore converts aggression to a relatively flat line (Fig. 4d; 4g, right). Thus, the superficially different courtship vs. aggression phenotypes caused by silencing pCd neurons can be explained by a common mechanism, whereby inhibition of pCd neurons increases the leak rate constant of a neural integrator, which may control a state of social arousal or drive^20, 41^.

### pCd neurons display neural integrator properties

We next investigated whether pCd neurons display integrator properties at the level of their physiology. The observation that they exhibit stepwise summation of P1 input (Fig. 1e, ED Fig. 3a) is consistent with this idea. Surprisingly, repeated direct stimulation of PPF1 neurons did not exhibit such summation, and evoked faster-decaying responses (median τ∼13.4 s) than evoked by indirect P1 activation (median τ∼83 s), indicating that persistent activity cannot be triggered cell autonomously (Fig. 5a). However, pCd function might be necessary, although not sufficient, for persistent activity (Fig. 5b, right). If so, then persistent pCd activity should not recover from transient inhibition performed during the decay phase following P1 stimulation^42, 43^. Alternatively, if pCd cells simply “inherit” persistence passively from an upstream input (Fig. 5b, left), their persistent P1 response should recover following transient inhibition. We therefore stimulated P1 neurons (5 s) whilst imaging from pCd cells, and after a short delay (25 s) briefly (∼10 s) inhibited pCd activity using the green light-sensitive inhibitory opsin GtACR1^44^ and 2-photon spiral scanning^45^ at 1070 nm to restrict inhibition to pCd cells (Fig. 5e, f and Methods).

Actuation of GtACR1 in pCd neurons following P1 stimulation caused a rapid, ∼68% decrease in ΔF/F signal, which did not recover to control levels following the offset of inhibition, but rather remained flat (Fig. 5g2, blue shaded area, solid vs. dashed line and Fig. 5h, pCd, green bar). This effect is not due to irreversible damage to pCd neurons by photo-inhibition, since reactivation of P1 neurons following transient pCd inhibition reliably re-evoked pCd persistent activity, and multiple cycles of P1 stimulation with or without GtACR1 actuation could be performed with consistent results (ED Fig. 7a, b, pCd). Furthermore, 2-photon spiral scanning at 1070 nm of pCd neurons lacking GtACR1 had no effect (Fig. 5g3), confirming that the decrease in GCaMP signal is due to inhibition of activity by GtACR1 and not to 2-photon irradiation. As the experiment was originally performed using Fru-LexA to label pCd cells, we confirmed the result using a pCd-specific driver (ED Fig. 8, blue shading).

As an additional control, we also performed the same manipulation on PPF2 neurons, another FruM^+^ population located near pCd (Fig. 5c), which also showed persistent responses to P1 activation (Fig. 1b3; Fig. 5d, PPF2). In this case, following GtACR inhibition PPF2 activity quickly recovered to the level observed at the equivalent time-point in controls without 1070 nm photo-inhibition (Fig. 5g5, 5h, PPF2 and ED Fig. 7b, PPF2). Thus, PPF2 activity is not required continuously to maintain a persistent response to P1 activation. In contrast, persistence in pCd neurons requires their continuous activity. However the fact that persistent activity cannot be evoked by direct stimulation of pCd neurons alone suggests that persistence likely requires co-activation of a network comprised of multiple neurons.

### pCd neurons are required for an effect of females to persistently enhance male aggressiveness, and are activated by an aggression-promoting pheromone, cVA

The foregoing data indicated that pCd neurons are required to maintain a P1 activation-triggered persistent internal state, which prolongs wing extension in solitary males and promotes aggression when male flies encounter another male. We next asked whether pCd neurons are similarly required for a persistent internal state triggered by naturalistic cues. Since P1 neurons are activated by female cues (reviewed in ref. ^14^), we examined the influence of transient female exposure on male aggressive behavior. Previous studies have demonstrated that females can enhance inter-male aggression (ref^36, 46^ and see Fig. 3h vs. k), but whether this effect can persist following the removal of females was not clear. To investigate this, we pre-incubated individual male flies for 5 min with or without a live female, and then gently transferred them into an agarose-covered arena to measure their aggression (Fig. 6a). Male flies pre-incubated with a female showed significantly higher levels of lunging than controls (Fig. 6b), indicating a persistent influence of female exposure to enhance aggressiveness.

We next asked whether this persistent influence requires continuous pCd activity. To do this, male flies expressing GtACR1 in pCd neurons were pre-incubated with females, and briefly photostimulated with green light during the aggression test (Fig. 6c). Transient inhibition of pCd neurons abrogated the effect of female pre-exposure to enhance aggression (Fig. 6d), mirroring the effect of such transient inhibition to disrupt persistent physiological activity in these cells (Fig. 5g2). Thus, continuous pCd neuron activity is required to maintain a persistent behavioral state change induced by female presentation. Importantly, this effect was not observed when P1 neurons were transiently silenced using GtACR, although such silencing of P1 cells did transiently disrupt male courtship towards females (ED Fig. 9), as previously reported^17^.

The foregoing experiments indicated that when males are removed from the presence of females and confronted with another male, their behavior switches from courtship to aggression. To investigate whether pCd neurons themselves might also play a role in the detection of male cues that trigger this behavioral switch, we investigated whether they can respond to 11-cis Vaccenyl Acetate (cVA), a male-specific pheromone that has been shown to promote aggression^47^ (Fig. 6e). Notably, cVA has already been shown to activate pCd cells in females^27^, where the pheromone promotes sexual receptivity. Although other pheromones have been shown to promote male aggression in *Drosophila*, such as 7-tricosene^48^, the non-volatility of that compound made it difficult to deliver in a controlled manner to walking flies in our imaging preparation (Fig. 6f) without physically disturbing them.

To do this, we imaged pCd activity using GCaMP6s in flies exposed to the following stimuli at 5 minute intervals: 10 s of P1 activation; cVA vapour presentation; or P1 stimulation (10 s) followed 30 s later by cVA (Fig. 6g1-3). Among pCd neurons persistently activated by P1 stimulation (Fig. 6g1), only half responded to cVA alone (defined as >2σ above baseline; Fig. 6g2). However, delivery of cVA 30 s after P1 stimulation (i.e., during the persistent phase of the response) yielded cVA responses (>2σ above post-P1 activity) in 90% of the pCd cells (Fig. 6g3). Moreover, peak cVA responses were significantly greater following P1 activation, than in flies exposed to the pheromone on its own (median increase 1.8-fold; Fig. 6h, i). Thus, individual pCd neurons that are activated by P1 stimulation in males can also respond to cVA (Fig. 6e), and this response is enhanced during the persistent phase of the P1 response.

## DISCUSSION

Optogenetic activation of P1 neurons evokes both courtship song, in a reflexive manner^15, 16^, and a persistent internal state of arousal or drive^20^ that promotes aggression in the presence of a conspecific male^9, 35^. Here we have identified a population of indirect persistent P1 follower cells, called pCd neurons^27^, whose activity is necessary for P1-triggered persistent aggression. pCd neurons are also necessary for persistent UWEs triggered by P1 activation, on a time scale outlasting P1 activity (as measured in separate imaging experiments). An earlier study^17^ reported that P1 activity is continuously required during male courtship following initial female contact, but did not distinguish whether this requirement reflected continuous stimulation of P1 cells by non-contact-dependent female-derived cues (e.g., motion cues^11, 14^), or a true fly-intrinsic persistent response. In contrast, the use of transient optogenetic stimulation here clearly demonstrates persistent fly-intrinsic responses. Nevertheless we cannot exclude that persistent P1 activity may occur during natural courtship bouts^17^. Importantly, however, we show that pCd but not P1 neurons are required for a persistent increase in aggressive state induced by transient female pre-exposure (Fig. 6d). Together, these data suggest that pCd neurons participate in a network that may encode a persistent memory of a female, which can be combined with the detection of an opponent male at a later time to elicit aggression^9, 20^. The observation that P1 neuron activation enhances pCd responses to cVA, an aggression-promoting pheromone^47^, is consistent with this idea.

Our physiological data suggest that pCd neurons are part of a circuit that temporally integrates P1 input to yield a slow response that decays over minutes (Fig. 1e). The fact that transiently silencing pCd neurons using GtACR irreversibly interrupts this slow response argues that it indeed reflects persistent pCd activity, and not simply persistence of GCaMP6s fluorescence. It is likely that this integrator circuit comprises additional neurons, including non-Fru-expressing neurons.

Evidently, P1 neurons activate this circuit in parallel with a “command” network, including pIP10 descending interneurons^7, 49^, that triggers rapid-onset courtship behavior. These results illustrate how acute and enduring responses to sensory cues may be segregated into parallel neural pathways, allowing behavioral control on different time scales, with different degrees of flexibility (Fig. 6j). The incorporation of parallel neural pathways that allow behavioral responses to stimuli to be processed on multiple timescales may represent an important step in the evolution of behavior, from simple stimulus-response reflexes to more integrative, malleable responses^41, 50, 51^.

Our data raise several new and interesting questions for future investigation. First, what cells provide direct synaptic inputs to pCd neurons, and what is the connectional relationship of these cells to P1 neurons? Second, the fact that pCd activity is necessary but not sufficient to trigger persistence suggests that other cells likely contribute to the integrator circuit; what are these cells (Fig. 6j, Y, Z)? Finally, how is persistence encoded, and what is the role of pCd neurons in determining its duration? The data presented here provide insight into the complex networks that underlie behavioral temporal dynamics^17, 52^ in *Drosophila*, and offer a useful point-of-entry to this fascinating problem.

## Acknowledgements

We thank H. Inagaki for comments on the manuscript, B. Pfeiffer and G. M. Rubin for fly strains, A. M. Wong for help with initial development of all-optical stimulation and imaging, A. Sanchez for maintaining fly stocks, C. Chiu for lab management and G. Mancuso for administrative assistance. This work was supported in part by NIH grant R01 DA031389. D.J.A. is an Investigator of the Howard Hughes Medical Institute. The authors declare no conflicts-of-interest.

## Author Contributions

D.A. and Y.J. conceived the project, designed experiments and co-wrote the manuscript; Y.J. performed all experiments, collected and analyzed data and prepared figures; A.K. performed mathematical modeling studies; H.C. generated R41A01-AD, R41A01-DBD, and nuclear-localized GCaMP6s constructs and flies; M.F. and A.C.-C. provided unpublished LexAop-GtACR flies.

## Data Availability

Datasets generated during the current study are available from the corresponding author on reasonable request.

## METHODS

### Rearing conditions

Flies were reared under standard conditions at 25°C and 55% humidity, on a 12 h light/12 h dark cycle. 2-5 days old virgin females were used to cross with different male stocks. The density of experimental flies (−5 pupae/cm^2^) was controlled by limiting the number of parents; crosses with too high or too low density of progeny were discarded. Male flies were collected 0-2 days after eclosion and reared either individually (single-housed) or at 18 flies (group-housed) per vial for 5-6 days before the behavioral assays. Newly eclosed males were excluded from collection. For optogenetic experiments, eclosed males were reared in the dark with food containing 0.4 mM all-*trans*-retinal (Sigma-Aldrich, St. Louis, MO). For two-color optogenetic experiments, flies were reared in the dark from larval stage. Virgin females provided during behavioral tests were reared at high density (30 flies per vial) for 2-3 days. Flies carrying Gal4 and UAS-opsin transgenes were maintained in the dark to prevent uncontrolled activation of the opsins.

### Fly strains

The following lines were generated in this study. *R41A01-LexA (vk00027* and *attp2)*, *R41A01-AD (attp40)*, and *R41A01-DBD (attp2)* were constructed based on the methods described in ref^53^. R41A01 enhancer fragment was amplified from genomic DNA based on sequences in (ref^54^). The primers used for amplification were designed based on recommendations in the Janelia FlyLight project and Bloominton Drosophila Stock Center (https://bdsc.indiana.edu/stocks/gal4/gal4_janelia.html). For making *LexAop2-NLS-GCaMP6s (su(Hw)attp5)*, two nuclear localization signal (NLS) peptides, one from SV40 and the other from the *Drosophila* gene *scalloped,* were used. SV40-NLS (ccaaagaagaaaaggaaggta) was fused to the 5’ end, and the scalloped-NLS (agaaccaggaagcaagtcagttcgcacatccaagtgctggctcgccgtaaactccgcgagatc) was fused to the 3’ end of the codon-optimized GCaMP6s. A DNA fragment containing syn21-SV40-NLS-GCaMP6s-scalloped-NLS was ligated into pJFRC19-13LexAop2-IVS-myr::GFP-sv40 (Addgene plasmid # 26224) via XhoI and XbaI restriction enzyme sites. The sv40 terminator in the pJFRC19 was replaced with p10 terminator via XbaI and FseI sites. To generate LexAop-GtACR1 flies, the GtACR1 Drosophila-codon-optimized sequence^44^ was subcloned into pJFRC19-13LexAop2-IVS-myr::GFP-sv40 (Addgene plasmid # 26224) plasmid. The GtACR1::eYFP fragment was swapped with the myr::GFP fragment using XhoI and Xba1. *15A01-LexA (attp2)*, *BDP-AD (attp40)* and *BDP-DBD (attp2)*, *10xUAS-NLS-tdTomato (VK00040)*, *13xLexAop2-NLS-GFP (VK00040)*, *10xUAS-Chrimson::tdTomato (su(Hw)attp1* and *attp18)*, *20XUAS-Chrimson::tdTomato (su(Hw)attp5)*, *13xLexAop2-myr::tdTomato (attp18)*, *13xLexAop2-OpGCaMP6s (su(Hw)attp8)*, *20xUAS-OpGCaMP6s (su(Hw)attp5), 13xLexAop2-mPA-GFP (su(Hw)attp8)*, *13xLexAop2-Kir2.1::eGFP (VK00027)*, *10xUAS-Kir2.1::eGFP (attp2)*, *10xUAS-GFP (attp2)*, *R21D06-LexA (attp2)* were from G. Rubin; *dsx-DBD* was from S. Goodwin^55^; *Fru-LexA* was from B. Baker^22^; *Orco-LexA* was from T. Lee^56^ ; *UAS-CD4::spGFP1-10* and *LexAop-CD4::spGFP11* were from K. Scott^57^; *20XUAS-GtACR1::eYFP (attp2)* was from A. Claridge-Chang; Wild-type Canton S was from M. Heisenberg^58^.

The following lines were obtained from the Bloomington Stock Center: *BDP-LexA (attp40)* (77691), *71G01-Gal4 (attp2)* (39599), *71G01-DBD (attp2)* (69507), *15A01-Gal4 (attp2)* (48670), *15A01-AD (attp40)* (68837), *R41A01-Gal4 (attp2)* (39425), *R41A01-LexA (attp40)* (54787), *R21D06-DBD (attp2)* (69873), *ChAT-DBD* (60318), *VGlut-DBD* (60313), *Gad1-p65AD* (60322), *UAS-Denmark; UAS-Syt-eGFP* (33064), *GH146-Gal4* (30026), *13XLexAop2-CsChrimson::mVenus (attp40)* (55138), *10XUAS-myr::GFP* (su(Hw)attp8) (32196), *10XUAS-myr::GFP* (attp2) (32197), *UAS-Kir2.1::eGFP* (6595).

### Two-photon GCaMP imaging

Calcium imaging was performed using a custom-modified Ultima two-photon laser scanning microscope (Bruker). The primary beam path was equipped with galvanometers driving a Chameleon Ultra II Ti:Sapphire laser (Coherent) and used for GCaMP imaging (920 nm). The secondary beam path was equipped with separate set of galvanometers driving a Fidelity-2 Fiber Oscillator laser (Coherent) for GtACR1 actuation (1070 nm). The two lasers were combined using 1030 nm short-pass filter (Bruker). GCaMP emission was detected with photomultiplier-tube (Hamamatsu). Images were acquired with an Olympus 40x, 0.8 numerical aperture objective (LUMPLFLN) equipped with high-speed piezo-z (Bruker). All images acquisition was performed using PrairieView Software (Version 5.3). For fast volume imaging (Fig. 1a, b and ED Fig. 1), three 4-µm optical sections were collected at 180 × 180 pixel resolution with a frame rate ∼0.83 Hz. All of the other images were acquired at 256 × 256 pixel resolution with a frame rate 1 Hz. Saline (108 mM NaCl, 5 mM KCl, 4 mM NaHCO3, 1 mM NaH2PO4, 5 mM trehalose, 10 mM sucrose, 5 mM HEPES, 0.5 mM CaCl2, 2 mM MgCl2, pH=7.5) was used to bathe the brain during functional imaging. Saline containing 90 mM KCl was added for high-resolution z stack after functional imaging to verify cell identity in ED Figure 1.

To prepare flies in vivo imaging, 6-8 days old flies were anesthetized on a cold plate and mounted on a thin plastic plate with wax. The wings, all legs, antenna, and arista were kept intact, wax-free, and free to move. Saline was added on the top side of the plate to submerge the fly head. A hole in the posterior-dorsal side of the head was opened using sharp forceps. Animals were then placed beneath the objective, and a plastic ball supported with air was positioned under the fly. The conditions inside of the imaging setup were maintained similar to the rearing conditions (25°C and 55% humidity). The flies were habituated for 30 min, and their behaviors were observed from the side using Point Grey Flea3 camera mounted with 0.5x-at-94 mm Infinistix lens fitted with a bandpass IR filter (830 nm, Edmund Optics) to block the two photon imaging laser and optogenetic stimulation lights. Animals that exhibited no movement, strenuous movement, and prolonged abdomen bending during and after habituation were discarded.

Chrimson activation during calcium imaging was performed as described in ref^15^. A deep red (660 nm) fiber-coupled LED (Thorlab) with band-pass filter (660 nm, Edmund Optics) was used for light source to activate Chrimson. A 200 µm core multimode optic fiber placed 200 µm away from the brain was used to deliver 10 Hz, 10 ms pulse-width light. The light intensity at the tip of optic fiber was set to be 39.2 µW. For two photon GtACR1 actuation, 1070 nm laser (Fidelity-2, Coherent) was delivered by galvanometers to a circular area with diameter =∼15 µm containing 1-3 cell bodies in focus for −10 s by spiral scanning (10 µm/pixel, 45.24 ms/repeat, 220 repeats). Galvanometers were re-calibrated weekly using a slide glass coated with thin layer of fluorescent dye. Field of view was adjusted in order to keep the spiral scanning area near the center of the imaging field. cVA was presented by directing a continuous airstream (80 mL/min) through a 4 mm diameter Teflon tube directed at the fly’s antennae. A custom-designed solenoid valve controller system was used to redirect the airstream between a blank cartridge and one containing cVA or Ethanol (solvent control). To make odour cartridges, 10 µL of undiluted cVA (Cayman Chemicals, 20 mg/mL) or Ethanol were placed on filter papers, and dried for 3 min to remove solvent before inserted into 15 mL pre-cleaned vials (Sigma-Aldrich).

### Imaging data analysis

All data analysis was performed in MATLAB (MathWorks). ROIs (region of interest) corresponding to individual cell bodies were manually selected and fluorescence signal from the ROIs were smoothed with a moving average (window =5 frames). For volume imaging (Fig. 1a, b and ED Fig. 1), a single focal plane in which we observed the highest ΔF/F was used for each cell. Normalized ΔF/F values for each trials were calculated by dividing ΔF/F by the maximum ΔF/F. The average signal before photostimulation was used as F0 to calculate the ΔF/F, and cells with peak ΔF/F responses < 4σ above baseline more than 1/3 trials were excluded. Decay constants (tau) were fit to minimize mean-squared error between observed ΔF/F traces and a five-parameter model of cell responses to optogenetic stimulation. Specifically, the ΔF/F trace evoked by three consecutive pulses of optogenetic stimulation was fit with a weighted sum of three impulse responses sharing a characteristic rise time tau_R and decay time tau: fit values of tau_R and tau were the same for all three evoked responses, while response amplitudes were fit independently. Fit impulse responses in the model were set to be 30 s apart, following experimental stimulation conditions. The best-fit 80% of cells (MSE<2.06) were used to generate plots of population-average responses. “Percent of peak” in Fig. 5h and ED Fig. 8c were calculated from mean normalized ΔF/F values between 10-30 s after GtACR1 actuation. cVA responses for Fig. 6h were calculated by subtracting mean GCaMP signal 10 s before cVA presentation from those obtained during cVA presentation (10 s). cVA responses from each cell delivered 30s after P1 stimulation were divided by cVA responses without concurrent P1 stimulation (cVA only), to calculate fold change (Fig. 6i). cVA alone or P1+cVA stimulation were delivered in random order following initial selection for P1-responsive pCd neurons. Individual cell responses used in Fig. 6g-i were the average of 2-3 trials per cell.

### Labeling neurons with Photoactivation after GCaMP imaging

Photoactivation experiments were performed in vivo using spiral scanning as described above. To perform GCaMP imaging and PA-GFP activation simultaneously, two Chameleon Ultra II Ti:Sapphire lasers (Coherent), one set at 920 nm and the other at 710 nm, are combined using 760 nm long pass filter (Bruker). Cell bodies of pCd neurons were identified by functional imaging using NLS-GCaMP6s, and a three-dimensional region of photoactivation was defined. The defined region of photoactivation was photoactivated by two cycles of spiral scanning (diameter =∼7.5 µm, 45.24 ms/repeat, 20 repeats, 150 ms inter-repeat-intervals) separated by 20 min interval to allow diffusion of photoactivated PA-GFP molecules to the projections. 20 min after second cycle of the spiral scanning, 3-dimensional images were acquired at 1024 × 1024 pixel resolution. To reduce the fly’s movement and residual GCaMP signal, cold saline containing 1mM EDTA was perfused until the end of image acquisition. tdTomato signals and photoactivated PA-GFP signals were imaged simultaneously at 940 nm. Non-PPF1 PA-GFP and NLS-GCaMP basal fluorescence have been masked for clarity and z stack were created (Fig. 1c3 and c4) using Fluorender^59^ and Fiji^60, 61^ software.

### Immunohistochemistry

Brains from 7-to-10-day-old adult files were dissected and stained as previously described^35^. The primary antibody mixture consisted of 1:1000 rabbit anti-GFP (Thermo Fisher Scientific, Cat#A11122), 1:1000 chicken anti-GFP (Aves Lab, Cat#GFP-1010), 1:100 mono-clonal (for GRASP experiment, ED Fig. 4j-r) mouse anti-GFP (Sigma-Aldrich, Cat#G6539), 1:1000 rabbit anti-DsRed (Takara Bio, Cat#632496), 1:50 mouse anti-Brochpilot nc82 (Developmental Studies Hybridoma Bank), and 10% normal goat serum (Sigma-Aldrich) in PBST. Secondary antibodies used were 1:1000 goat anti-rabbit-Alexa488 (Thermo Fisher Scientific, Cat#A11008), 1:1000 goat anti-chicken-Alexa488 (Thermo Fisher Scientific, Cat#A11039), 1:1000 goat anti-mouse-Alexa488 (Thermo Fisher Scientific, Cat#A11001), 1:1000 goat anti-rabbit-Alexa568 (Thermo Fisher Scientific, Cat#A11011), and 1:1000 goat anti-mouse-Alexa633 (Thermo Fisher Scientific, Cat#A21050).

Confocal stacks were obtained with Fluoview FV1000 or FV3000 (Olympus). Fiji^60, 61^ and Fluorender^59^ software was used to create z stack images. For brain registration (ED Fig. 4g-i), the two images shown in ED Fig. 4b and d are registered to T1 template brain^62^ using CMTK registration tools^63^.

### Behavioral assay

Temperature and humidity of the room for behavioral assay was set to 25°C and 55%, respectively. All naturally occurring behavior assays were performed between 2:00pm to 7:00pm. Optogenetically-induced behaviors were not performed at specific times. All the behavior assays except mating assay (ED Fig. 6) were performed in 8-well acrylic chamber (16 mm diameter x 10 mm height, modified from ref^15^, and side of the each well was coated with aInsect-a-Slip (Bioquip Products). Temperature probe (Vktech) was inserted into one side of the chamber to accurately monitor the chamber temperature. The clear top plates were coated with Sigmacote (Sigma-Aldrich), and the floor of the arenas was composed of clear acrylic covered with food (2.5% (w/v) sucrose and 2.25% (w/v) agarose in apple juice). Flies were introduced into the chambers by gentle aspiration using a mouth pipette, and the chambers were placed under the behavioral setup. Flies were allowed to acclimate to the chamber under the camera without disturbance for 90 s before the recording. Fly behaviors were recorded at 30 Hz using Point Grey Flea3 camera mounted with Fujinon lens (HF35HA-1B) fitted with a long pass IR filter (780 nm, Midwest Optical Systems). Camera was located ∼0.5 m above the chamber, and IR backlighting (855 nm, SmartVision Lights) was used for illumination from beneath the arena.

Optogenetic activation was performed as described previously^15^. Briefly, a 655 nm 10 mm Square LED (Luxeon Star) was used to deliver 0.48 mW/mm^2^ light for 30 seconds. For dead female presentation (Fig. 3d-f and j-l, Fig. 4a, and ED Fig. 9), 2-5 day old wild-type Canton S virgin females were freeze-killed, and affixed in the middle of the arena with UV curable glue. The ventral end of the female abdomen was glued to prevent copulation.

For the female induced aggression assay (Fig. 6a-d), single-housed male flies were transferred individually into empty vials containing a virgin female, and allowed to freely interact with the female for ∼5 min. After this pre-exposure period, the male flies were gently transferred to the behavior arena covered with 2.25% (w/v) agarose in dH2O, instead of fly food. For GtACR1 stimulation (Fig. 6c-d and ED Fig. 9), a 530 nm 10 mm Square LED (Luxeon Star) was used to deliver 117 μW/mm^2^ light for 10 seconds. Male flies that initiated copulation during the 5 min pre-exposure period were not tested.

For the mating assay (ED Fig. 6),12-well two-layer chambers in which the layers were separated by a removable aluminum film. 2-5 day old wild-type Canton S virgin females were introduced into the lower layers, and males of a particular genotype were introduced in the upper layers. Flies were allowed to acclimate to the chamber for 90 s as described above before removing film. Behavior recording started right after film was removed.

### Behavioral data analysis

Analysis of lunging and unilateral wing extension was performed as described in ref^9^. Briefly, fly posture was tracked from recorded videos using Caltech FlyTracker software, which is available for download at http://www.vision.caltech.edu/Tools/FlyTracker/, and bouts of behaviors were automatically annotated using the Janelia Automatic Animal Behavior Annotator (JAABA)^64^. All annotations were manually validated to remove false positives. Behavioral assays with dead females (Fig. 3d-f and j-l) were manually scored without using JAABA due to inaccuracy. Data shown in Fig. 3a-c and g-i were also manually scored for consistency. Copulation latency for ED Fig. 6 was manually scored, and the total number of males that had engaged in copulation was summed across the 30-min period and plotted as a percentage of total flies for each time point. Courtship bouts shown in ED Fig. 9 were manually annotated following the definition of courtship bouts described previously^17^. Statistical analyses were performed using Matlab and Prism6 (GraphPad Software). All data were analyzed with nonparametric tests. The cutoff for significance was set as an α<0.05. Each experiment was repeated at least twice on independent group of flies. Outliers were defined as data points falling outside 1.5x the interquartile range of the data, and were excluded from plots for clarity, but not from statistical analyses.

### Curve Fitting for Leaky bucket model

Rasters of courtship and lunging behavior in a 15-minute window were averaged across flies and binned in 10-second (for courtship) or 20-second (for lunging) time windows to produce a time-evolving population average behavior rate. Behavior rates for courtship and lunging were each fit with a three-parameter leaky integrator model with dynamics 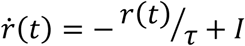, which has analytical solution 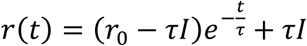, where *r* is the behavior rate as a function of time *tt* (in minutes), *II* is a constant sensory input, *ττ* is the time constant of integration, and *rr*_0_ is the initial behavior rate at the start of recording.

Parameters *II*, *ττ*, and *rr*_0_ were fit to minimize the mean squared error between model and data, for courtship and for lunging. Parameter values were jointly fit across the two behaviors (courtship and lunging) and across the four experimental conditions: pCd > Kir2.1 (manipulation), pCd > GFP, BPD > Kir2.1, and BPD > GFP (controls). To reduce the number of free parameters, the sensory input *II* was constrained to take the same value for all groups and conditions, while *rr*_0_ was fit separately for courtship and for aggression; only *ττ* was fit independently for each group and each behavior.

---

**Extended Data Figure 1.**
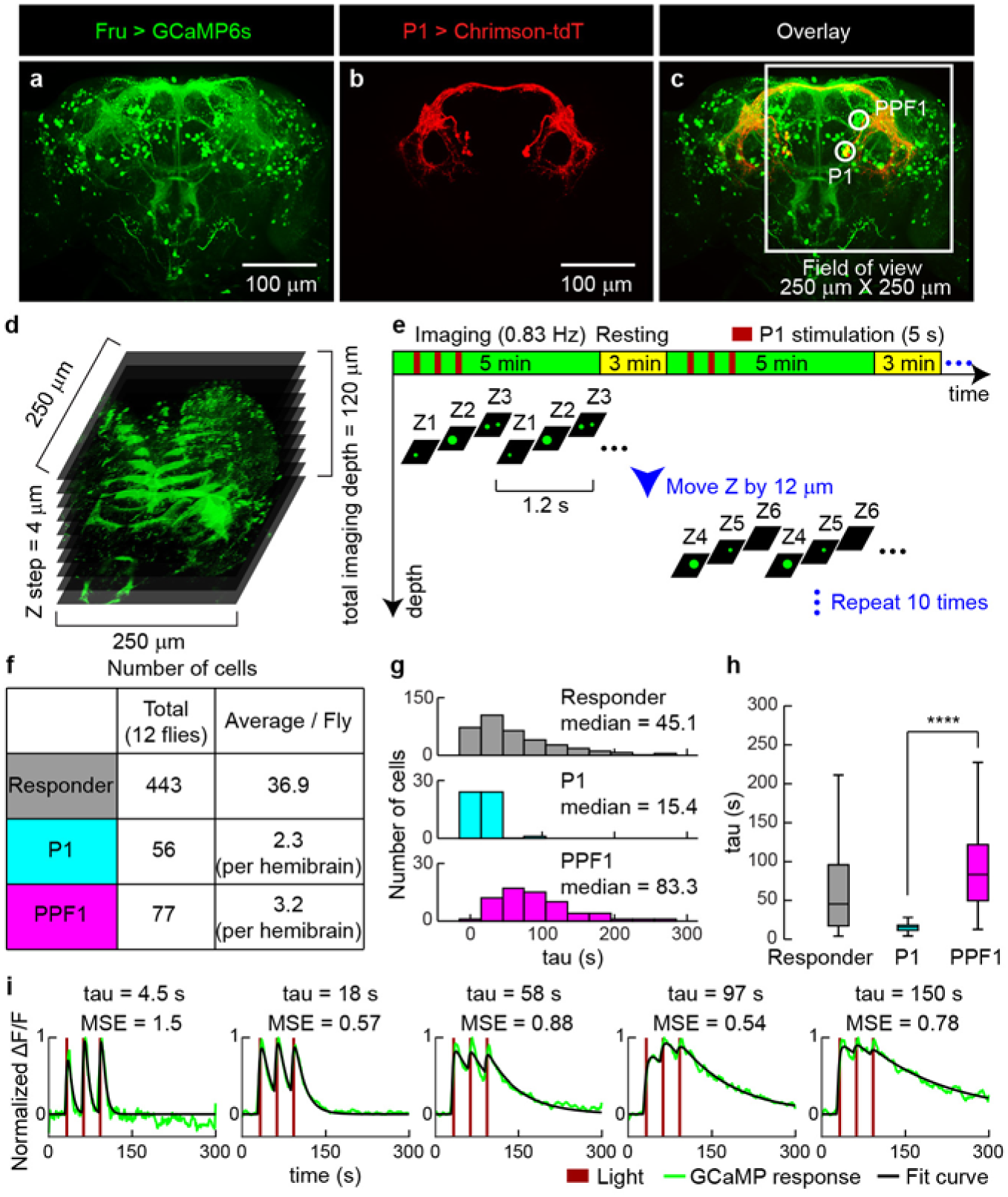
Volumetric functional GCaMP imaging to identify persistent P1 follower cells. (a-c) Maximum intensity confocal stacks showing projection patterns of Fruitless (a) and P1^a^ (b) neurons^9, 20^ expressing GCaMP6s and Chrimson-tdT, respectively; (c), overlay. (d-e) Schematics illustrating functional connectomics strategy. Responses to P1^a^ photostimulation (3 x 5 s pulses) from multiple Fru>GCaMP6s cells in each imaging plane (250 x 250 µm^2^) were recorded during ten 5 min trials, at multiple z-depths (4 µm/z-step) covering 120 µm. (f) Number of Fruitless^+^ cells that responded to P1^a^ activation. PPF1 cells were identified anatomically in high-resolution images acquired following P1 stimulation trials, using 40 mM KCl-containing saline to increase baseline GCaMP6s signals. Red channel (Chrimson-tdTomato) was used to identify P1 neurons, and cell body position and primary projection pattern were used to identify PPF1 neurons. P1 and PPF1 were visible in both hemi-brains of all specimens, but some responder cells on the lateral side appeared only in one hemi-brain (see Field of view marked in (c)). (g) Histogram of τ (tau, decay constant of a model exponential fit to observed neural ΔF/F traces) for all responder cells (top, grey), P1 cells (middle, light blue), and PPF1 neurons (bottom, magenta). (h) Quantification and statistical test for τ. Statistical test used was a Mann-Whitney U-test. **** *P* < 0.0001. τ from 80% of the total identified cells (MSE ≤ 2.06, 354 cells) were used for the plot (g) and quantification and statistical test (h). (i) Representative examples of GCaMP responses and τ for different responder cells. Dark red lines indicate Chrimson activation at 660 nm (3 stimulations, 5 s each, 10 Hz, 10 ms pulse-width, 25 s inter-stimulation interval).

**Extended Data Figure 2.**
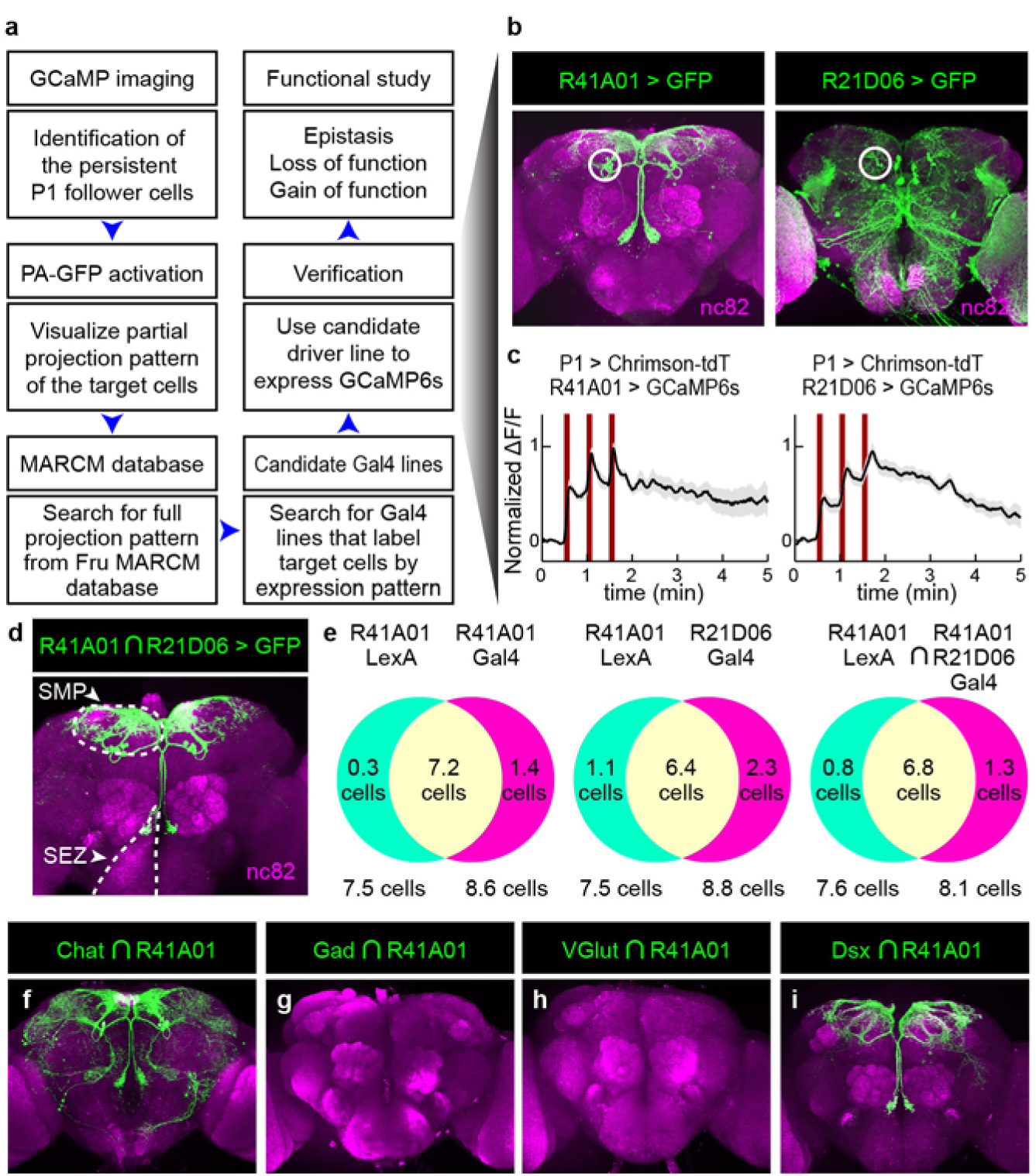
Gaining genetic access to PPF1 neurons and molecular phenotype of pCd neurons. (a) Flowchart of the protocol for identifying specific Gal4 lines labeling PPF1 neurons. (b) Anatomy of two Gal4 lines, R41A01 (left) and R21D06 (right) that label PPF1 neurons. Maximum-intensity projection (z-stack) of confocal 2-µm optical sections. (c) Functional imaging of putative PPF1 neuronal cell bodies labeled by R41A01-LexA (left) and R21D06-LexA (right). Traces represent normalized ΔF/F response to P1 stimulation (dark red bars, 3 repeats of 5 s stimulation, 10 Hz, 10 ms pulse-width, 25 s inter-stimulation interval), and were obtained from cell bodies within the white circles indicated in (b). Mean±sem, n=7 (4 flies) for R41A01, and 9 (4 flies) for R21D06. (d) Anatomy of split-Gal4 intersection between R41A01-AD and R21D06-DBD in the male brain. SMP and SEZ are indicated with white dashed line. (e) Quantification of pCd cell numbers (per hemibrain) labeled by two different reporters, UAS>tdTomato and LexAop>GFP, in flies co-expressing the indicated GAL4 or LexA drivers. Green=GFP positive, Red=tdTomato positive, Yellow=double positive. Area of Venn diagram not scaled to number of cells. n=12 hemibrains per test. (f-i) Anatomy of split intersection between R41A01-AD and Chat-DBD^66^ (f), Gad1-AD and R41A01-DBD (g), R41A01-AD and VGlut-DBD (h), and R41A01-AD and dsx-DBD. Maximum-intensity projection of confocal 2-µm optical sections.

**Extended data Figure 3.**
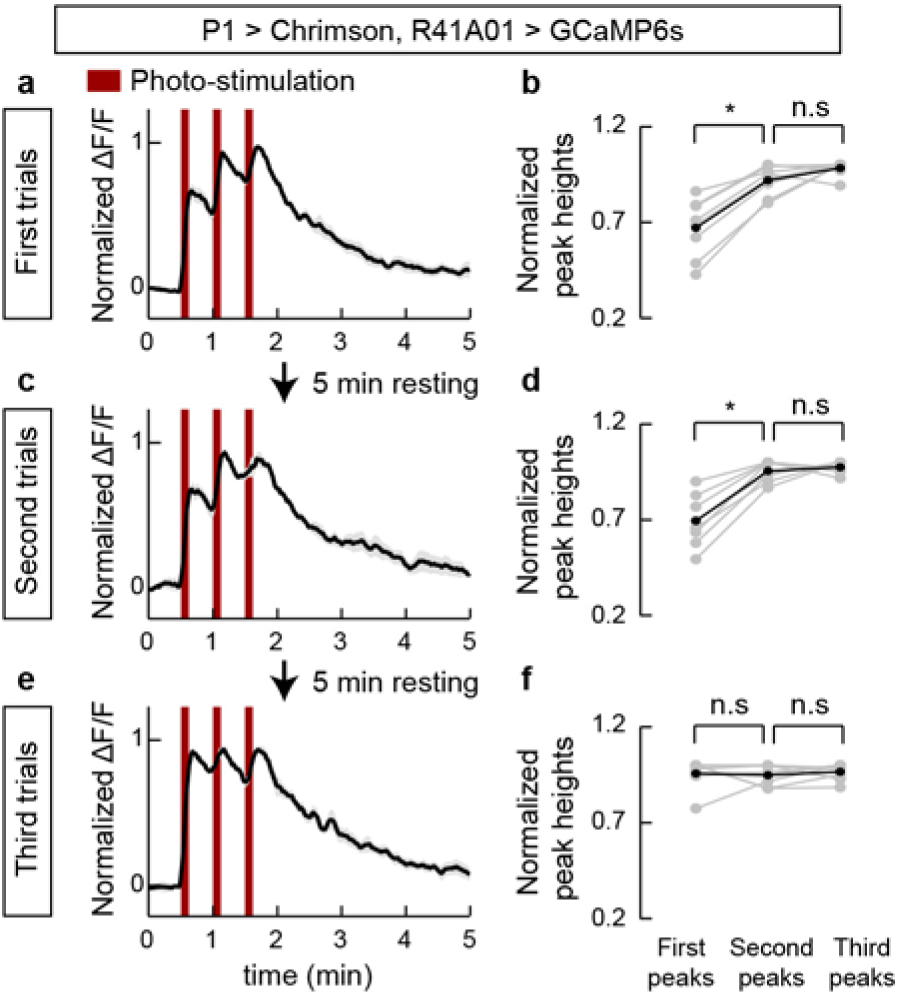
Integration of repeated P1 input by pCd neurons. (a, c, e) Normalized GCaMP response of pCd neurons to optogenetic stimulation of P1 neurons. Mean±sem. n=8 cells, 6 flies. (b, d, f) Normalized peak heights during each P1 stimulation. Statistical test used was Wilcoxon signed test with correction for multiple comparisons. * *P* < 0.05

**Extended Data Figure 4.**
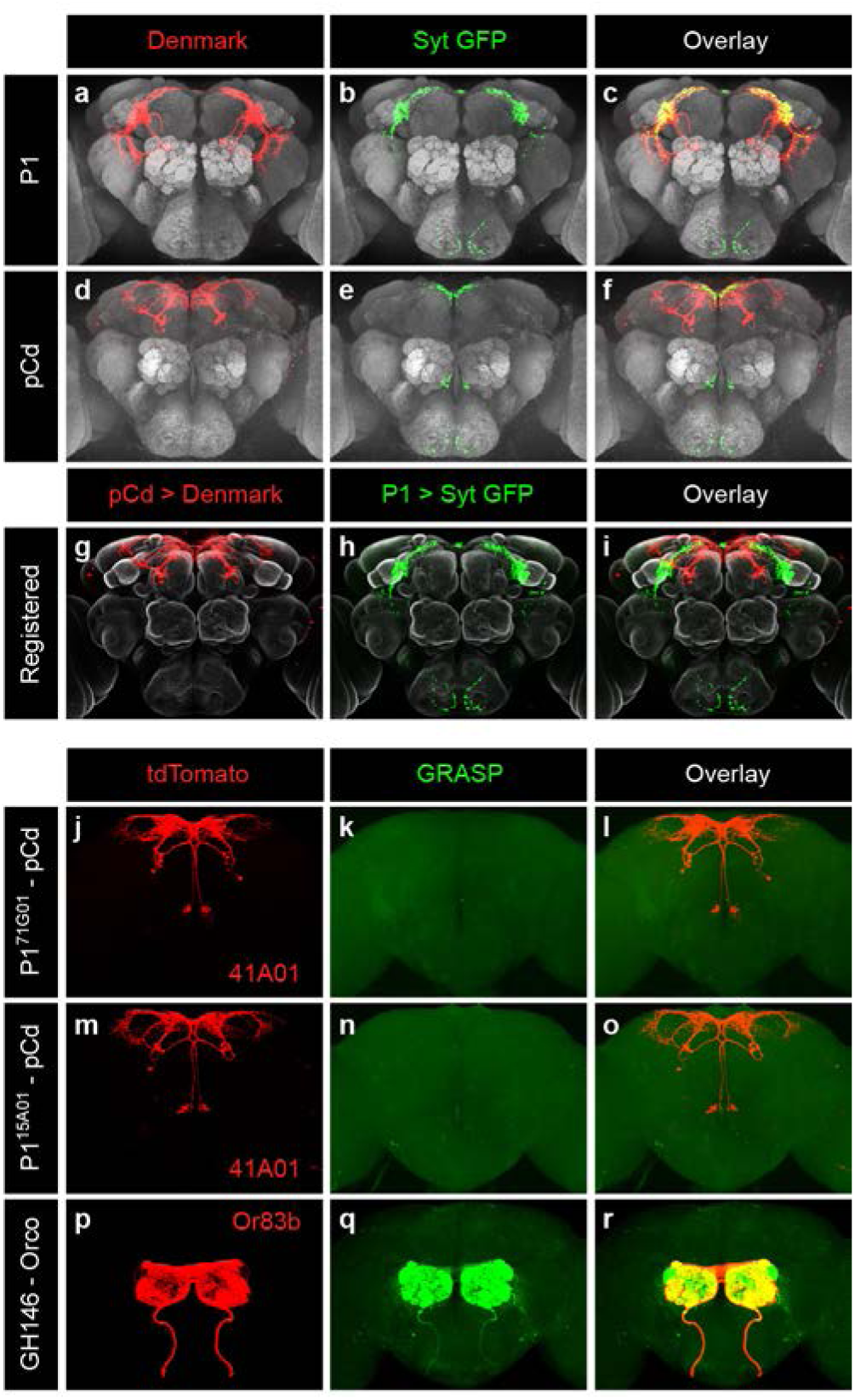
Anatomic relationship between P1 and pCd neurons. (a-f) Input and output region of the P1 and pCd neurons visualized by double-labeling with somatodendritic marker (Denmark, red) and pre-synaptic marker (Syt-GFP, green). (g-i). Co-registered images showing somatodendritic region of pCd neurons and pre-synaptic region of P1 neurons. Note that yellow regions in (i, “Overlay”) are not observed when the image is rotated and viewed from a different angle, indicating a lack of overlap. (j-r) GRASP^34^ experiments performed between R41A01 (pCd driver) and either of two P1 drivers, 71G01 (j-l) and 15A01 (m-o), or between GH146 and Orco as a positive control (p-r). tdTomato was expressed in one of the putative synaptic partners, R41A01 (j and m) or Orco (p), to mark fibers for detailed analysis. No positive GRASP signal is observed between pCd and either of the 2 P1 drivers (j-o).

**Extended Data Figure 5.**
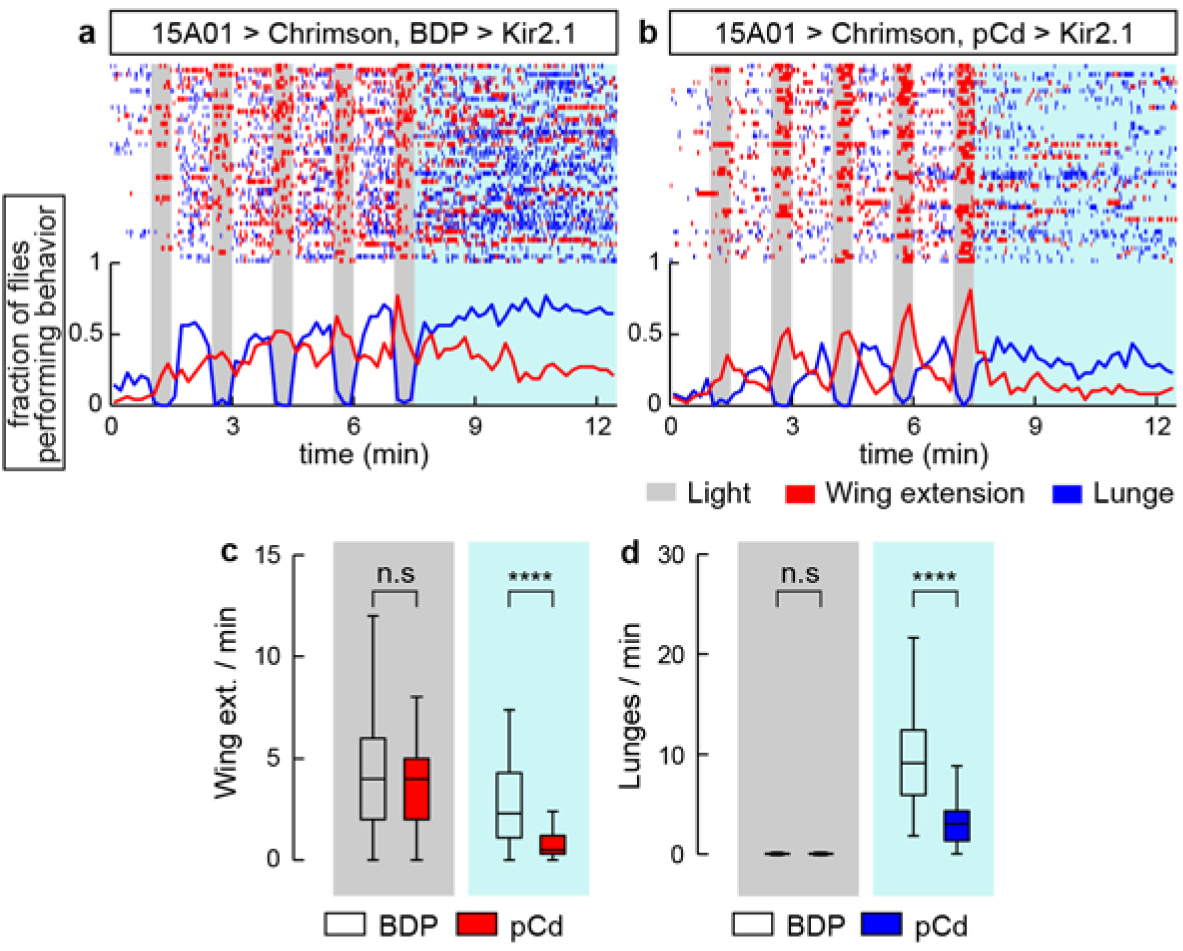
Inhibition of pCd neurons with R41A01∩R21D06 Split-Gal4 reduces P1-induced social behaviors. (a-b) Top: raster plot showing wing extensions (red ticks), and lunges (blue ticks) in pair of males. In this experiment, a single driver 15A01-LexA^9, 35^ was used to activate P1 neurons, while a split GAL4 driver (ED Fig. 2d) was used to inhibit pCd neurons, complementing the genetic strategy used in Fig. 2 in which a split-Gal4 was used to activate P1 neurons, while R41A01-LexA was used to inhibit pCd neurons (see Table 1 for genotypes). Bottom: fraction of flies performing unilateral wing extensions (red lines), and lunges (blue lines) in 10 s time bins. Gray bars indicate Chrimson activation (5 repeats of 30 s stimulation, continuous light, 60 s inter-stimulation interval). n= 48 flies per genotype. (c-d) Quantification and statistical tests for unilateral wing extensions (c) and lunges (d) during P1 stimulation (gray shading) and after photostimulation (blue shading), without (open boxes, BDP) or with (red boxes) silencing of pCd neurons using Kir2.1. **** *P* < 0.0001 for between-genotype comparisons (Mann-Whitney U-test). Note that both wing-extensions and aggression are suppressed by pCd silencing during the post-P1 stimulation period.

**Extended Data Figure 6.**
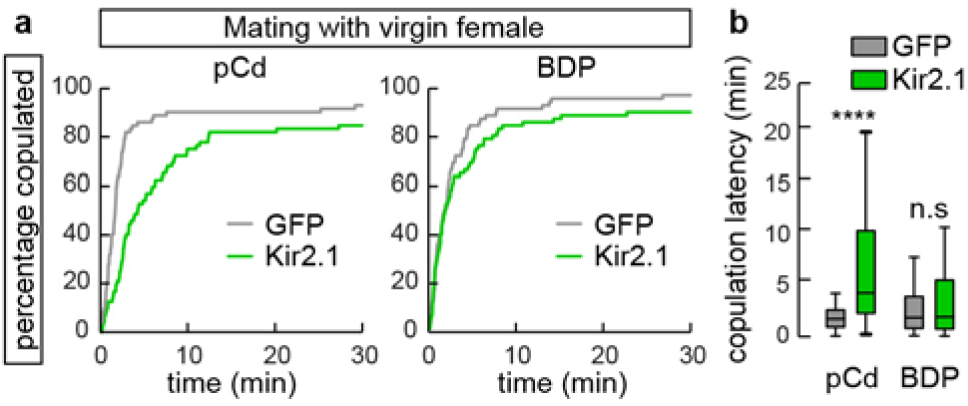
Inhibition of pCd neurons increases copulation latency. (a) Individual males of the indicated genotypes were paired with a live wild-type virgin female. Cumulative percentage of flies that copulated over 30 min is shown. (b) Quantification and statistical tests for copulation latency. **** *P* < 0.0001 for between-genotype comparisons (Mann-Whitney U-test).

**Extended Data Figure 7.**
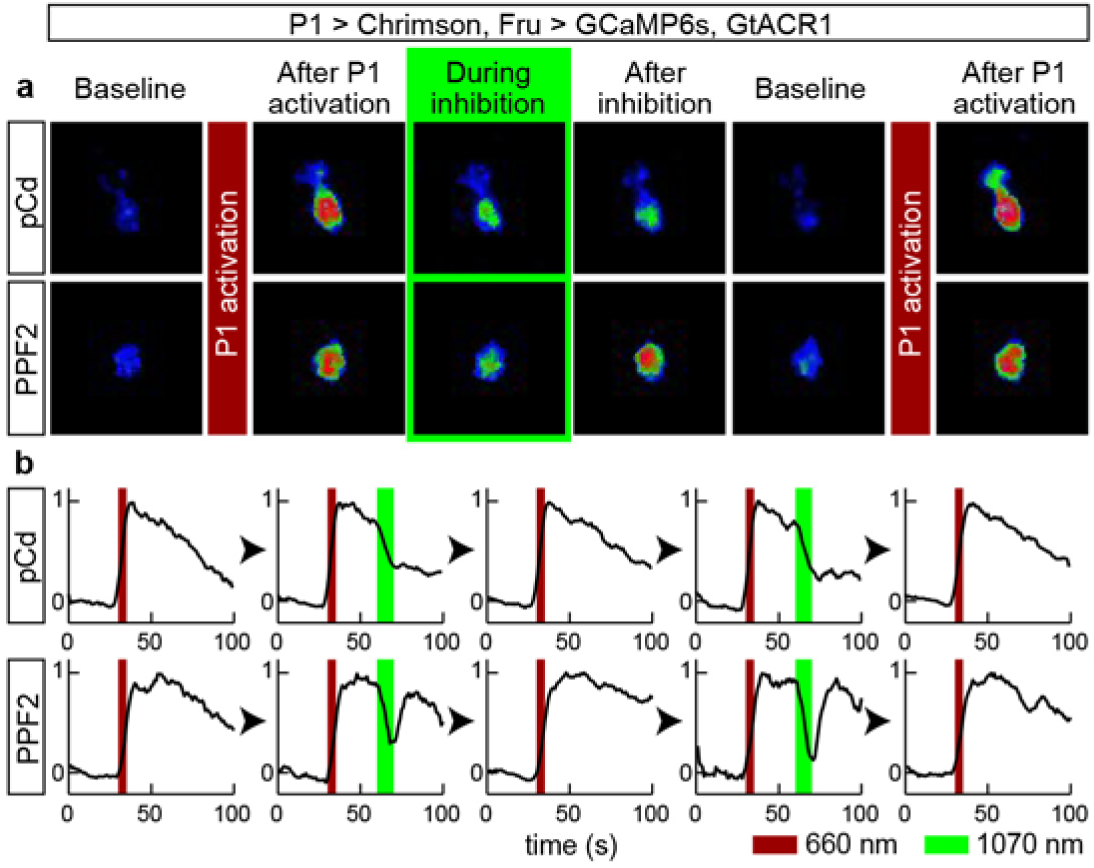
Multiple cycles of P1 stimulation and GtACR1 actuation in pCd and PPF2 neurons. (a) Representative GCaMP fluorescent images of pCd (*upper*) and PPF2 neurons (*lower*) at different time points following Chrimson-mediated P1 stimulation (wide-field LED actuation at 660 nm), and cell-restricted GtACR-mediated pCd or PPF2 inhibition (2-photon spiral scanning actuation at 1070 nm). pCd and PPF2 neurons both respond to P1 stimulation, and their response endures following offset of P1 photostimulation (“After P1 activation”). GCaMP signals in pCd neurons rapidly decrease upon photo-inhibition (“During inhibition”, green outline), and do not recover 10 s following offset of GtACR actuation (“After inhibition”). In contrast, PPF2 activity recovers after photo-inhibition. pCd and PPF2 neurons were reliably reactivated by a second cycle of P1 stimulation after following GtACR-mediated inhibition. Images shown are averaged over 5 frames. (b) Representative GCaMP trace (normalized ΔF/F) from individual trials. Multiple cycles of P1 stimulation with or without GtACR1 actuation did not change the initial responses of pCd and PPF2 neurons to P1 stimulation. Dark red bar indicates Chrimson activation at 660 nm (5 s, 10 Hz, 10 ms pulse-width), and green bar indicates GtACR1 actuation (∼10 s, spiral scanning) 25 s after Chrimson activation.

**Extended Data Figure 8.**
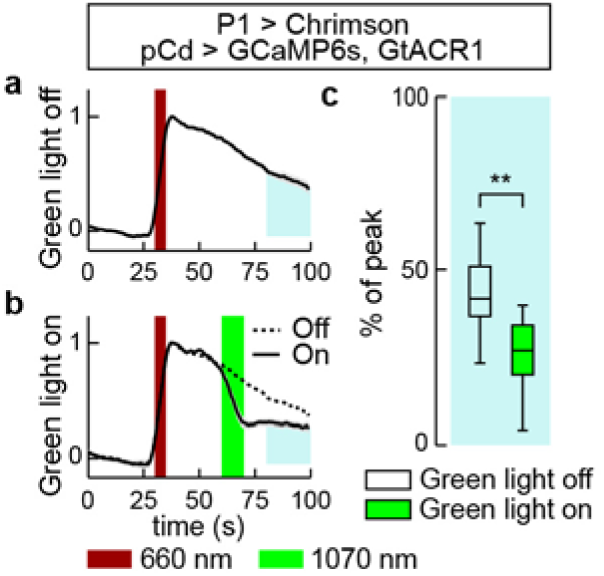
GtACR-mediated inhibition of pCd neurons following P1 stimulation labeled by a pCd-specific driver. (a) GCaMP6s response of pCd neurons (normalized ΔF/F) labeled with the driver R41A01-LexA (pCd^R41A01^) to P1 stimulation (dark red bar) without GtACR1 actuation. (b) GCaMP6s response of pCd^R41A01^ to P1 stimulation with GtACR1 actuation. n=10 trials from 3 flies (a-b). Dark red bar indicates Chrimson activation at 660 nm (5 s, 10 Hz, 10 ms pulse-width), and green bar indicates GtACR1 actuation (∼10 s, spiral scanning) 25 s after Chrimson activation. (c) Normalized area under the curve after photo-inhibition (blue shaded area in (a-b)). Statistical test used was a Mann-Whitney U-test. ** *P* < 0.01. This experiment confirms the result reported in Fig. 6, in which Fru-LexA was used to express GCaMP6s and pCd neurons were identified morphologically

**Extended Data Figure 9.**
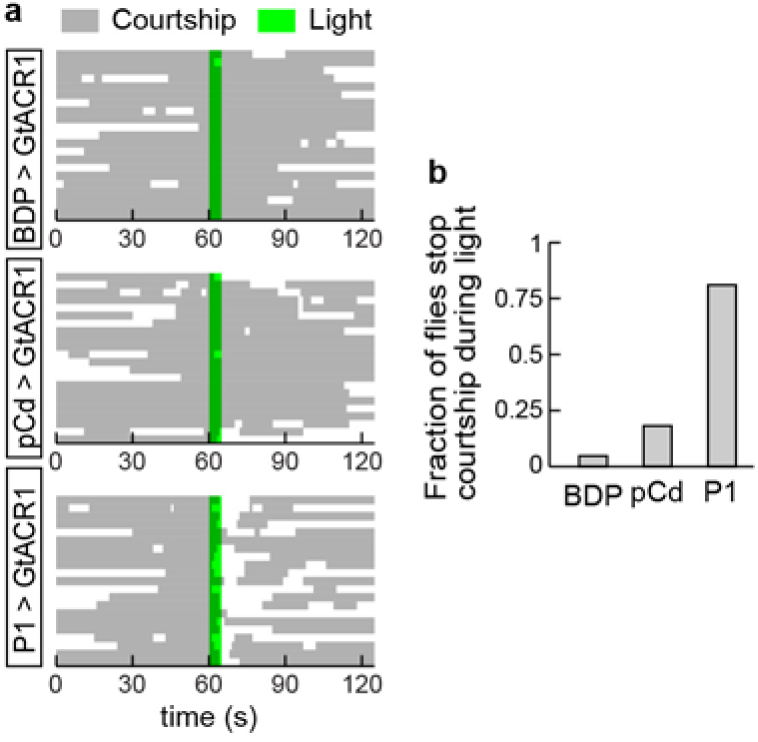
Transient inhibition of P1 neurons interrupt ongoing courtship behavior toward dead female. (a) Raster plot showing courtship toward dead female (gray). Note that “courtship” metric used here incorporates multiple behavioral actions, following the definition used by Zhang et al.^17^, and thereby differs from the wing extension metric used in other figures (see Methods for details). Green line indicates GtACR1 stimulation (530 nm, 10 Hz, 10 ms pulse-width) for 10 s. n=21 for BDP and P1 > GtACR1, and 22 for pCd > GtACR1. (b) Fraction of flies stop on-going courtship behaviors during light stimulation.

